# Carbon and microbes in a degrading palsa mire are distinct from peatland and a wider connected sub-Arctic fluvial system

**DOI:** 10.1101/2025.05.07.652394

**Authors:** Nea Tuomela, Samu Elovaara, Jenni Hultman, Hermanni Kaartokallio, David N. Thomas

## Abstract

Climate change is altering the biogeochemical cycling of carbon and nutrients in the northern peatland and permafrost regions, which provide one of the largest terrestrial carbon storages. Lateral transfer of carbon needs to be more widely studied, especially in smaller streams and catchments, as they receive high loading of organic matter and are hotspots of carbon degradation. In this study we combined measurements of dissolved organic matter (DOM) quality and quantity with microbial community data from a small Arctic catchment. Our aim was to understand how the catchment is affected by two sub-catchments, degrading palsa permafrost mire and peatland thawing in spring. The small thaw ponds in the palsa mire were clearly distinct from the rest of the catchment and ponds in the peatland: Palsa ponds had higher DOM concentration, more aromatic DOM and distinctive microbial communities compared to the peatland ponds and the rest of the catchment. DOC export rates from the palsa and peat sites were comparable at the time of sampling, but local DOM processing was higher in the palsa site. We also detected high abundances of ultra-small Patescibacteria. Patescibacteria dominated the microbial community composition in all the sampled waters.

## Introduction

Northern peatlands and permafrost are two of the largest reservoirs of organic carbon on Earth, storing a total of 415 Pg C, of which 185 Pg of carbon is in permafrost peatlands [1]. The Arctic is undergoing rapid warming [2,3], which is leading to the mobilization and release of soil organic carbon from permafrost [4]. Palsas, isolated peat mounds with a permafrost core, are especially vulnerable to increasing temperatures and changing rainfall patterns [5] and drastic degradation of palsas has been detected pan-Arctic [6–8]. Following permafrost degradation, organic matter and nutrients can be released into ponds and streams, to be transported and transformed [1,9–11]. Fluxes of dissolved organic matter (DOM) are highly seasonal, with highest export rates during spring freshet [12]. However, with permafrost thaw changing the hydrology, DOM export patterns and quality can change [11,13,14].

The properties of DOM along with temperature and nutrient availability determine whether it is degraded or transferred further downstream [15]. In general DOM with lower molecular weight and less aromatic structure is more available for microbial degradation and consumed rapidly in lower-order streams [16]. Ancient permafrost DOM has been characterised as being highly available to microbial degradation, due to containing material comprised of DOM with lower molecular weight and lower aromaticity [10,17,18]. However, in other studies permafrost derived DOM has been characterized as poor substrate for microbial consumption [13,19]. Regardless of the quality of organic matter released from soils to freshwaters, the increasing runoff of organic matter affects the function of fluvial networks in the Arctic region [13,20]. The role of headwaters and thaw ponds in processing permafrost DOM is pertinent because they receive high loading of DOM from soils [15,21,22].

Microbial communities and their functionality are one of the key factors in DOM processing [23,24] and vice versa, DOM quantity and quality is an important driver for microbial community assembly [25–27]. Microbial assemblages in headwaters are also highly affected by the input of microbial taxa from soil [27–31]. Of the soil microbial taxa, some will increase in abundance downstream from the soils, acting as seed taxa for the downstream communities [32]. The community formation processes change from stochastic in the headwaters towards deterministic further downstream [30,32].

Headwater streams and ponds have been studied to an extent, but there is a knowledge gap on microbial community composition and function [15,33]. For example, less known phylum Candidatus Patescibacteria can represent most of the free-living microbial communities in surface and groundwaters [34–37]. Patescibacteria have been understudied due to their ultrasmall size, leading to underestimation of their abundance with standard water sampling protocols [35,38,39]. Patescibacteria might possess genes involved in the biogeochemical cycling of carbon and nitrogen [38] and degrading peatland and permafrost derived organic matter to usable forms for methanogens [40].

We examined DOM quality and quantity, and microbial communities in a small scale sub-arctic river catchment. We collected samples along the catchment, from small ponds and headwater streams to larger riverine sites. We then compared the results from the palsa mire to the non-permafrost peatland on the same catchment. We hypothesized that the high DOM input from degrading palsa soils would be visible all the way downstream by distinct DOM composition and microbial community assembly. We expected DOC concentration to be higher, and the DOM quality to differ in palsa mire compared to peatland site. We also expected the microbial communities to reflect the differences in DOM quantity and quality.

## Materials and Methods

### Sampling site and sampling strategy

Samples were collected in June 2023 from Kidisjoki River catchment, a sub-catchment of the Utsjoki River close to Kevo Subarctic Research Institute (69° 48’ N, 27° 06.5’ E) in Finnish Lapland (Figure 1). The Kidisjoki River has a pristine catchment, with the only direct human impact being caused by a reindeer herding facility (operational during October-November). The location has subarctic climate with cumulative rainfall in of 7.5 mm and average temperature of 11.9 °C in July 2023 [41]. Kidisjoki catchment area covers 22 km² of which approximately 1 km² is palsa permafrost mire [42]. We sampled along the edges of two of the closest palsa mounds in the mire, which were surrounded by thaw ponds. Palsas were characterized by collapse features along the edges indicating degradation of the mound [43] (Supplementary Figure 1). The vegetation of Kidisjoki River catchment consists mainly of mountain birch, with treeless alpine heaths, peatland and Scots pine present in the lower parts of the basin. A more detailed description of the vegetation can be found in [43,44].

**Figure 1.**
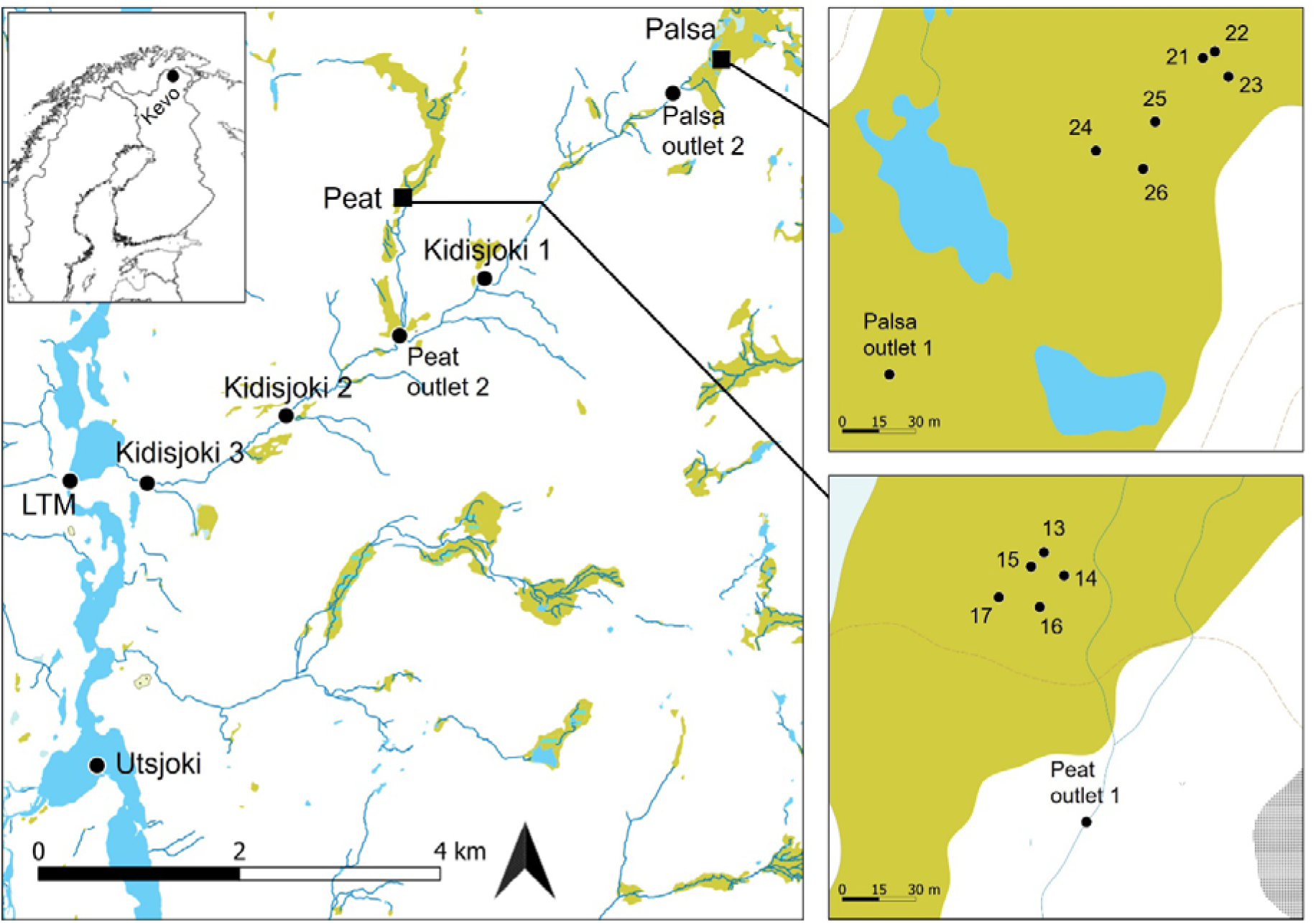
Map of the sampling scheme in Kidisjoki River catchment, Kevo, Finland. Sampling sites are indicated with black dots. Water samples were collected from all sites and accompanied with soil samples from sampled pond sites, from peatland labelled as 13-17 and from palsa site labelled as 21-26. Palsa samples were collected along the edges of two palsa mounds (21-23 and 24-26). Kidisjoki River was sampled from three locations (Kidisjoki 1-3). LTM stands for long term monitoring site. Green colour is indicative of peatland, and base map was acquired from National Land Survey of Finland [95].

Water samples were collected from Utsjoki River (n=1), along Kidisjoki River (n=3), from peatland streams (n=2) and ponds (n=5) and palsa streams (n=2) and ponds (n=6) (Figure 1). Water was collected into clean acid-washed plastic bottles and transported in coolers to the field station for further processing. When present, large particles were removed by pre-filtering the water with plankton net (sieve size 40 µm). Temperature and conductivity were measured with YSI 6600 V2 CTD sonde (YSI Corporation, Yellow Springs, Ohio, USA) and stream flow rates were measured with ADV FlowTracker (SonTek, San Diego, CA, USA).

All water samples for microbial communities were processed within the sampling day. Autoclaved filtration units (Millipore, 250ml) were used to sequentially filter the sample with 0.45 µm and 0.1 µm pore size polyether sulfone membrane filters (Sartorius, Göttingen, Germany). Samples were filtered until maximum of 30 minutes had passed to keep cells intact. Filtered volumes are listed in Supplementary Table 1. Filters were folded with sterile tweezers and stored in cryovials at -80 ^∘^C until DNA extraction.

Soil samples were collected next to the ponds from peatland and palsa sites (Figure 1). Six replicate soil profiles were collected from palsa site and five from peatland. Soil was collected from three depths. From peatland we sampled at 2, 4 and 10 cm below soil surface with a shovel. The peat layer was shallow, approximately 15 cm, on top of mineral soil and vegetation comprised mainly of sedges. Palsa soil was collected by hammering a steel pipe (inner diameter 3 cm) into the soil [43]. Samples were collected from three layers: surface layer (20-25 cm below soil surface), depth of freezing (23-33 cm) and frozen peat soil (49-59 cm) to total number of 18 samples. Samples were sliced with ethanol-disinfected knife, put into sterile plastic bags and stored in a cooler with -80 ^∘^C icepacks until transfer to a -80 ^∘^C freezer within hours.

### DOM degradation experiment

Microbial degradation of DOM was investigated by incubating water samples in the dark at 10 ^∘^C and collecting subsamples for analyses at the start and after approximately 15, 100, 150 and 220 days. The first sampling interval aimed to capture the short-term processes that could be related to the microbial community observed in situ. The subsequent samplings assessed the long-term resistance of DOM to general microbial degradation, as the microbial community was expected to change radically, and we thus did not attempt to connect this change to the observed components of the microbial community.

500 mL of sample was filtered through 0.8 µm to remove flagellate grazers, stored in a glass bottle and stirred periodically. In addition to filtered samples, unfiltered samples were collected at sites Palsa outlet 2, Peat outlet 2 and Main 3 to investigate how the presence of full planktonic community would affect degradation of DOM. Samples for the measurement of dissolved O_2_ were collected separately into glass stoppered vials with no head space and measured using Winkler titration to calculate microbial respiration.

Respiration was calculated only at the first subsampling interval. Optical properties of DOM were measured at each subsampling, bacterial abundance was measured at the first, second and the last subsampling and nutrient concentrations were measured at the first and the last subsampling.

### Microbial analysis

#### DNA extraction, sequencing and bacterial cell enumeration

DNA extractions for soil and water samples were done with DNeasy PowerSoil Pro Kit (Qiagen, Hilden, Germany). For soil samples, 0.25-0.30 g of soil was used for DNA extraction avoiding the exposed surface of the sample. For water samples, DNA was extracted from the filters and separately for the two size fractions. Filters were transferred to PowerBead tubes, Solution CD1 was added, and filter was broken with sterile pipette tip. Cell disruption step was done with FastPrep homogenizer (MP Biomedicals, Heidelberg, Germany; 30s, 5.5m/s). The rest of the extraction was completed following manufacturer’s instructions. DNA concentration was measured with Qubit Fluorometer, using Qubit High sensitivity dsDNA kit (Invitrogen by Thermo Fisher Scientific, Life Technologies Corporation, USA). DNA from the two size fractions was combined prior to sequencing. Libraries were prepared with Nextera™ DNA Flex Library Preparation Kit (Illumina, San Diego, CA, USA) and paired-end sequencing was done with AVITI (Element Biosciences, 2×150bp) at the Institute of Biotechnology, University of Helsinki. Bacterial cell abundance was determined using flow cytometry with LSRFortessa (BD Biosciences) at University of Helsinki Flow Cytometry Unit (Supplementary Methods 1).

#### Bioinformatics

Raw sequences were trimmed using cutadapt v4.9 [45] to remove Nextera adapters. Quality was checked with FastQC v0.11.9 [46] and MultiQC v1.19 [47]. After trimming, the sequences from three individual lanes were combined for each sample. Since our focus was in bacterial and archaeal communities, the 16S rRNA gene fraction of the sequenced metagenomes was analyzed. Trimmed reads were mapped against SILVA 138.1 database with PhyloFlash v3.4.2 [48]. Mitochondrial and chloroplast reads were removed prior to analysis with phyloseq package v.1.42.0 [49] in R v4.2.3 [50] and RStudio v2024.12.0 [51]. Bacterial and archaeal communities are presented as mean relative abundance for each sample type. Community compositions for individual samples can be found in Supplementary Figures 2-4.

### Water chemistry

Samples for dissolved organic carbon (DOC) and coloured dissolved organic matter (CDOM) were prepared by filtering 20mL of water through pre-combusted GF/F filters (4501^∘^C, 41h) into acid washed and pre-combusted glass vials. DOC samples were acidified to pH 2 with concentrated orthophosphoric acid, sealed with a septum cap and stored at −201^∘^C. DOC concentrations were analysed with TOC-L total organic carbon analyser (Shimadzu).

CDOM samples were stored in +4^∘^C until analysis after field campaign, within 10 days of sampling. Storage has been shown in previous studies to have negligible changes to CDOM and humic fluorescence characteristics of filtered samples [52–57]. Optical properties of DOM were conducted following [58]. CDOM absorption was measured with a Shimadzu 2401PC spectrophotometer using 1cm quartz cuvette over spectral length range 200-800 nm with 1 nm resolution, using ultrapure water (MQ) as blank. Absorption coefficient at 254 nm (a_CDOM(254)_) was used as a general indicator of optically active molecules and color of water. Ultraviolet specific absorbance at 254 nm (SUVA_254_) was calculated by normalizing DOC absorbance at 254 nm^-1^ to DOC concentration in mg L^-1^ [59]. CDOM slopes (S_275-295_, S_350-400_, S_300-650_) were calculated with R package cdom v0.1.0 [60].

Excitation–emission matrices (EEMs) of FDOM were measured with a Varian Cary Eclipse fluorometer (Agilent). The characteristics and the quality of the DOM were assessed with fluorescence peaks [61] extracted from the EEMs utilizing the R package eemR [62]. A blank sample was subtracted from the EEMs, and the Rayleigh and Raman scattering bands were removed from the spectra after calibration. EEMs were calibrated by normalizing to the area under the Raman water scatter peak 11 (excitation wavelength of 350 nm) of an MQ water sample run on the same session as the samples and were corrected for inner filter effects with absorbance spectra [63]. Fluorescence peaks T and C were used as proxies of protein-like and humic-like DOM, respectively [61]. In addition, the following indices were calculated: FI = fluorescence index [64], HIX = humification index [65], BIX = biological index [66]. We did not measure iron, which can be high in thaw ponds [21]. Iron has been detected to cause increase of on average 4-17% in SUVA_254_ values [67–69] and lower fluorescence signal by 7-23% due to quenching [68]. Therefore, we have only drawn conclusions based on large differences in CDOM variables, not on absolute values.

Nutrient concentrations (nitrate, nitrite, phosphorus, silicate and total dissolved nitrogen) were measured with SEAL AA500 AutoAnalyzer (Seal Analytical, US) using instrument specific applications of the standard colorimetric methods [70]. Ammonium concentrations were determined fluorometrically using Varian Cary Eclipse fluorometer (Agilent) [71].

### DOC export rates

Export rates of dissolved organic carbon (DOC) were calculated by multiplying the measured DOC concentration and flow rates for all our stream sites, except for the Utsjoki River. For the latter, DOC export rate was calculated by multiplying our DOC measurement with the flow rate of the sampling day from the daily flow rate measurement from the long-term monitoring site [72]. These DOC export rates were then scaled to a day specific export rates (kg C day^1^), assuming DOC concentration and flow rates remain the same.

### Soil properties

Soil physicochemical properties were measured from lyophilized and ground soils following [73]. Soil pH was measured following international standard ISO10390. Loss on ignition (LOI) was used to analyse soil organic matter (SOM) content according to Finnish standard SFS3008. Soil total carbon and total nitrogen contents were determined with a high temperature induction furnace (LECO, 828 series) [43].

### Statistical analysis

All data analysis were conducted in R v4.2.3 [50] and RStudio v2024.12.0 [51] with package tidyverse v2.0.0 [74]. Variation of biogeochemical properties was analysed with Principal Components Analysis (PCA, prcomp-command) utilizing R package ggbilot v0.55 [75]. Mean values of technical replicates were used and variables which were not determined for all samples were excluded. Screeplot was used to justify the selection of principal components (Supplementary Figure 7). For microbial community analysis phyloseq package v1.42.0 was used [49] and variation between samples was visualized with Principal Coordinate Analysis (PCoA) on genus level, utilizing Bray-Curtis distance. The microbial community taxonomic diversity was measured with Shannon Index (Supplementary Figure 6). Distance-based redundancy analysis was conducted with vegan package v2.6-4 [76] to assess how microbial community composition on genus level was affected by environmental properties. dbRDA was performed on selected environmental variables (DOC, slope ratio, SUVA_254_, hix, bix, fi, temperature, conductivity, pH, phosphate, nitrite, nitrate and total nitrogen), which were measured for all samples. Stepwise selection was used to only include significant predictors, which were hix, pH, phosphate and nitrite. Minimal collinearity (<3) was confirmed with variance inflation factor (VIF).

## Results

### Water chemistry

DOC concentrations were highest in palsa ponds, ranging from 2 992 to 9 330 µmol L^-1^, followed by peat ponds with values from 210 to 5 750 µmol L^-1^ (Table 1). Peat and palsa outlets had similar DOC concentrations of 407 and 401 µmol L^-1^ respectively. Kidisjoki River and Utsjoki River had similar range of DOC with each other. In peat ponds, a_CDOM(254)_ was relatively low compared to the DOC concentration, indicating possibly higher ratio of non-coloured DOM of the total DOM pool than in e.g. palsa ponds. The humification index was highest in palsa ponds, followed by peat outlet and the Kidisjoki River. Strikingly, the humification index decreased from palsa ponds to the palsa outlet, while the opposite was measured between the peat ponds and peat outlet. The lowest SUVA_254_ values were measured in peat ponds and the highest in palsa ponds. Palsa outlets had slightly higher SUVA_254_ than rest of the catchment. SUVA_254_ is a proxy for aromaticity [59], and the values would indicate higher aromaticity in palsa ponds than rest of the samples and the lowest aromaticity in peat ponds. Aromaticity increased from the peat ponds to the peat outlet and decreased from palsa ponds to the palsa outlet site.

**Table 1.** Physicochemical properties of freshwater sampling sites in the studied Kidisjoki River catchment in Utsjoki region, Finland. Number of sampling sites are expressed in column header.

| Parameter |  | Unit | Palsa pond<br>(6) | Palsa outlet<br>(2) | Peat pond<br>(5) | Peat outlet<br>(2) | Kidisjoki River<br>(3) | Utsjoki River<br>(1) |
| --- | --- | --- | --- | --- | --- | --- | --- | --- |
| DOC | | $\mu\text{mol L}^{-1}$ | 5,561 (2,217) | 407 (71.00) | 3,706 (2,882) | 401 (56) | 223 (5.9) | 237 |
| Organic N | | $\mu\text{mol L}^{-1}$ | 110.47 | 7.81 (2.04) | 4.30 (0.37) | 6.66 (0.65) | 3.80 (0.29) | 6.84 |
| DOC:DON |  |  | 46.13 | 53.37 (4.84) | 870.64<br>(709.65) | 59.91 (2.64) | 58.92 (4.93) | 34.69 |
| SUVA<sub>254</sub> | | $\text{L mg}^{-1} \text{m}^{-1}$ | 16.04 (3.99) | 8.41 (0.97) | 2.11 (2.69) | 7.98 (0.29) | 7.96 (0.06) | 7.19 |
| $\text{NO}_2^-$ | | $\mu\text{mol L}^{-1}$ | 1.02 (0.34) | 0.08 (0.01) | 0.04 (0.02) | 0.07 (0.01) | 0.08 (0.01) | 0.04 |
| $\text{NO}_3^-$ | | $\mu\text{mol L}^{-1}$ | 0.02 (0.04) | 0.02 (0.02) | 0.09 (0.03) | 0.08 (0.06) | 0.34 (0.34) | 0.13 |
| $\text{PO}_4^{3-}$ | | $\mu\text{mol L}^{-1}$ | 5.66 (4.37) | 0.03 (0.01) | 0.03 (0.01) | 0.04 (0.00) | 0.04 (0.00) | 0.03 |
| Si | | $\mu\text{mol L}^{-1}$ | 1.44 | 26.88 (16.28) | 60.21 (14.45) | 18.45 (5.24) | 91.85 (36.50) | 82.39 |
| TDN | | $\mu\text{mol L}^{-1}$ | 119.13 (38.02) | 7.95 (2.05) | 4.51 (0.34) | 6.88 (0.74) | 4.34 (0.60) | 7.11 |
| $\text{NH}_4^+$ | | $\mu\text{mol L}^{-1}$ | 2.13 | 0.04 (0.00) | 0.08 (0.00) | 0.06 (0.03) | 0.12 (0.03) | 0.11 |
| pH |  |  | 4.84 (0.99) | 6.25 (0.91) | 6.91 (0.16) | 7.19 (0.30) | 7.22 (0.18) | 5.57 |
| Conductivity | | $\mu\text{S cm}^{-1}$ | 95.15 (12.33) | 79.23 (1.77) | 79.83 (4.48) | 72.00 (2.00) | 77.00 (0.82) | 82.08 |
| Temperature | | $^{\circ}\text{C}$ | 14.89 (1.89) | 11.00 (2.78) | 13.25 (1.42) | 13.29 (1.43) | 6.80 (0.59) | 15.55 |
| ODO |  | % | 67.45 (15.52) | 75.25 (3.93) | 73.64 (10.80) | 89.50 (0.26) | 91.80 (0.25) | 99.46 |
| ODO | | $\text{mg L}^{-1}$ | 6.81 (1.51) | 8.35 (0.97) | 7.72 (1.16) | 9.38 (0.27) | 11.20 (0.15) | 9.91 |
| fi |  |  | 1.03 (0.10) | 1.10 (0.02) | 1.18 (0.08) | 1.16 (0.10) | 1.17 (0.03) | 1.04 |
| hix |  |  | 33.48 (6.80) | 23.64 (3.82) | 18.96 (7.70) | 37.97 (4.92) | 35.14 (5.69) | 17.43 |
| bix |  |  | 0.34 (0.03) | 0.51 (0.07) | 0.49 (0.07) | 0.41 (0.04) | 0.39 (0.02) | 0.42 |
| $S_R$ | | | 0.95 (0.07) | 0.75 (0.03) | 0.74 (0.02) | 0.72 (0.01) | 0.77 (0.01) | 0.86 |
| $S_{275-295}$ | | $10^{-3} \text{ nm}^{-1}$ | 8.88 (0.50) | 11.93 (0.13) | 12.92 (0.31) | 12.58 (0.29) | 13.00 (0.23) | 14.45 |
| $S_{350-400}$ | | $10^{-3} \text{ nm}^{-1}$ | 9.38 (1.04) | 16.03 (0.41) | 17.42 (0.14) | 17.50 (0.14) | 16.83 (0.41) | 16.73 |
| $S_{300-650}$ | | $10^{-3} \text{ nm}^{-1}$ | 18.23 (1.02) | 14.67 (0.14) | 15.36 (0.32) | 15.32 (0.25) | 15.48 (0.09) | 15.71 |
| Peak T |  | R.U. | 0.03 (0.00) | 0.08 (0.03) | 0.05 (0.03) | 0.04 (0.01) | 0.02 (0.00) | 0.04 |
| Peak A |  | R.U. | 0.45 (0.12) | 0.50 (0.01) | 0.34 (0.02) | 0.62 (0.06) | 0.34 (0.01) | 0.28 |
| Peak M |  | R.U. | 0.27 (0.08) | 0.31 (0.01) | 0.23 (0.01) | 0.39 (0.04) | 0.20 (0.00) | 0.19 |
| Peak C |  | R.U. | 0.31 (0.08) | 0.35 (0.02) | 0.24 (0.01) | 0.43 (0.04) | 0.24 (0.01) | 0.19 |
| $\alpha_{\text{CDOM}(254)}$ | | $\text{m}^{-1}$ | 981 (252.13) | 40.28 (2.41) | 19.83 (1.37) | 38.20 (3.99) | 21.28 (0.61) | 20.47 |
| HNA bacteria | | $10^6 \text{ cells ml}^{-1}$ | 5.74 (2.43) | 2.58 (0.57) | 2.39 (0.52) | 2.39 (0.76) | 1.22 (0.30) | 5.15 |
| LNA bacteria | | $10^6 \text{ cells ml}^{-1}$ | 4.73 (3.84) | 2.24 (0.78) | 1.75 (0.31) | 1.66 (0.59) | 0.95 (0.41) | 1.59 |

Total dissolved nitrogen (TDN) and phosphate (PO_4_ ^3-^) had highest concentrations in the palsa ponds. Concentration of TDN was 10-fold lower in other sampling sites. Palsa ponds had highest conductivity, followed by the Utsjoki River. Peat ponds, palsa outlet and the Kidisjoki River had similar conductivity, whereas peat outlet had slightly lower value. Oxygen concentration and saturation were lowest in palsa ponds, with slight increase to palsa outlets. Similar pattern was also observed in peat ponds and peat outlet. Bacterial numbers were highest in palsa ponds for both high and low nucleic acid bacteria, whereas palsa outlet and peatland had similar range of bacterial cells. Oxygen measurements and bacterial numbers indicated higher bacterial activity in palsa ponds than rest of the catchment. Palsa ponds had the highest temperature of all sites; they were dark coloured and warmed in the sunlight during the sampling day.

Principal component analysis of the nutrients, and DOM quality and quantity revealed distinction between the palsa pond sites and rest of the water samples (Figure 2). PC1 explained 44.7% and PC2 15.3% of the variation. PC1 variation was contributed by e.g. oxygen, spectral slopes, pH, SUVA_254_, conductivity, DOC, phosphate and nitrogen. The largest contributions to PC2 were Peak A, Peak C and Peak M. The samples were separated by PC1, with palsa ponds forming one group and the rest of the samples as another group. Palsa and peat outlets formed slightly into their own group.

**Figure 2.**
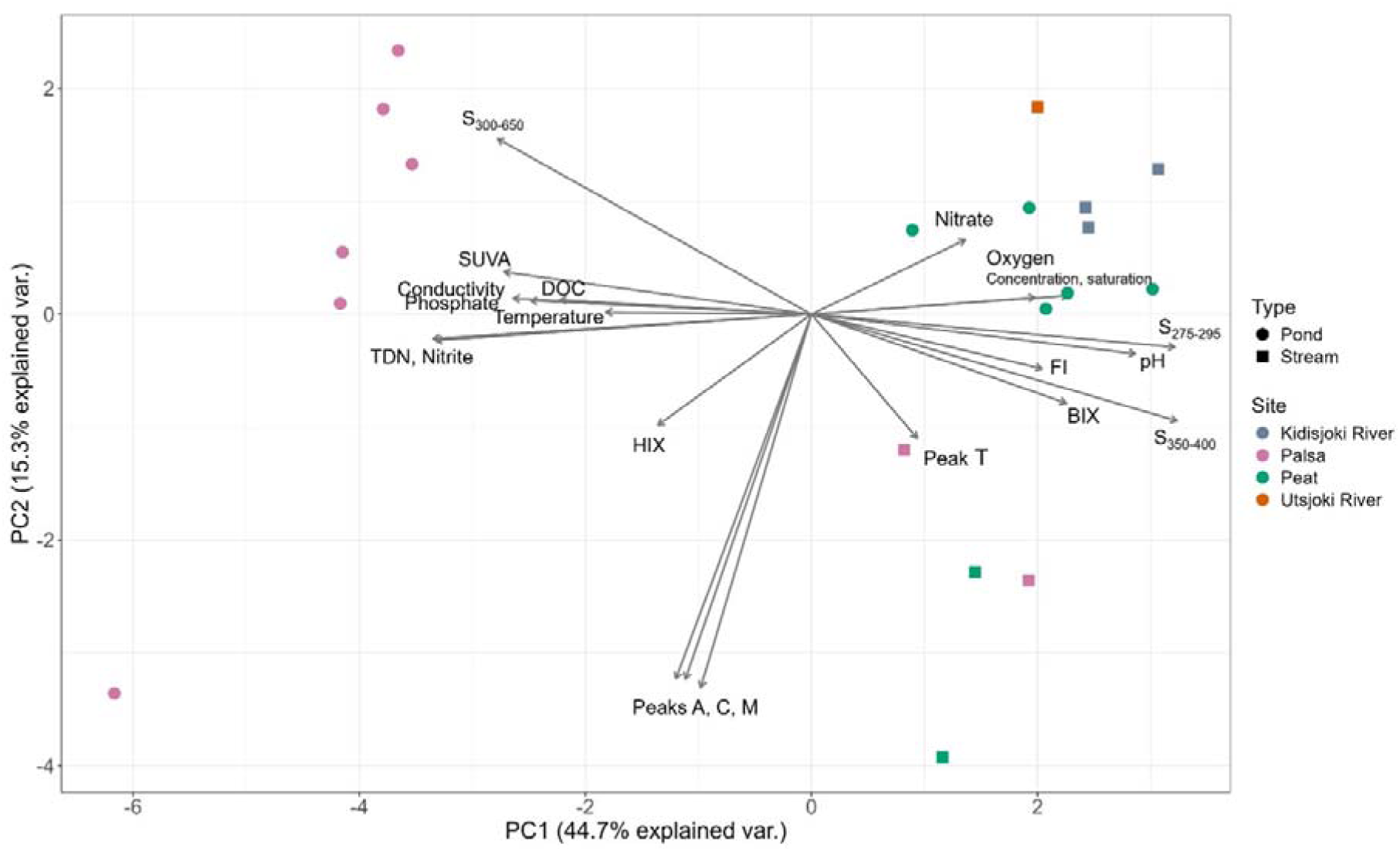
Principal component analysis of biogeochemical properties of the studied arctic catchment water samples from small ponds and streams. Measured variables include DOC concentration (DOC), nutrient concentrations and physicochemical properties. PCA includes also optical property measurements of DOM (Peaks A, C, M and T, Fluorescence index (FI), Biological index (BIX), Humification index (HIX), Spectral slopes (S_275-295_, S_350-400_ and S_300-650_) and SUVA (SUVA_254_). Sample type is indicated by shape and site by colour. Labels refer to the PCA loading arrows.

### Soil properties

The palsa soils were acidic with pH between 3.75 and 4.10. Palsa soils had C:N (total C : total N) ratios of between 35.2 and 38.7 and high SOM contents of 97% in all depths (Table 2). In peatlands the pH values were between 5.16 and 5.31, C:N ratio ranged from 17.81 to 25.85, and SOM was between 19 and 67%. SOM content in the peatland soils was highest in the soil surface.

**Table 2.** Soil properties in palsa and peatland soils, grouped by depth. Values are means of 6 replicates for each depth in palsa site and 5 for peatland samples. C:N ratio is the ratio of total carbon to total nitrogen. Standard deviation is given in parentheses. Palsa soil depths represent the dry ice free peat (20-25cm below soil surface), depth of freezing during the sampling (23-33cm) and frozen peat soil (49-59cm). Peat sample depths are expressed as the depth below soil surface.

| Site | Depth | pH | C:N ratio | Soil organic matter % |
| --- | --- | --- | --- | --- |
| Palsa<br>n = 6 | 20-25 cm | 3.75 (0.08) | 37.35 (7.71) | 96.98 (2.95) |
|  | 23-33 cm | 3.78 (0.10) | 35.20 (4.09) | 97.25 (1.27) |
|  | 49-59 cm | 4.10 (0.39) | 38.67 (9.50) | 97.93 (0.70) |
| Peat<br>n = 5 | 2 cm | 5.31 (0.22) | 25.85 (7.68) | 67.14 (28.37) |
|  | 4 cm | 5.19 (0.26) | 23.51 (4.50) | 59.68 (35.04) |
|  | 10 cm | 5.16 (0.15) | 17.81 (1.40) | 18.88 (19.88) |

### Microbial communities

#### Bacterial community composition

Based on the 16S rRNA gene annotations from metagenomes, the bacterial communities in the water samples were dominated by phyla Patescibacteria, Proteobacteria, Bacteroidetes, Acidobacteriota and Verrucomicrobiota (Figure 3a). Acidobacteriota, Actinobacteriota and Proteobacteria were abundant in palsa soil communities. In peat soils relative abundance was more evenly distributed among phyla, including e.g. Chloroflexi, and Verrucomicrobiota. Patescibacteria was a highly abundant phylum in palsa outlet, peat ponds, peat outlet and Kidisjoki River. In Utsjoki River, the community was dominated by Actinobacteriota, Bacteroidetes and Gammaproteobacteria. In order level the Utsjoki River bacterial community consisted of Burkholderiales, Frankiales, Flavobacteriales, Cytophagales and Chitinophagales (Figure 3b). Palsa pond community consisted of Burkholderiales, Acidobacteriales and Acetobacteriales. Palsa outlets, peat outlets, peat ponds and the Kidisjoki River were dominated by the orders Omnitrophales, Burkholderiales and from the Patescibacteria phylum, orders Candidatus Nomurabacteria and Candidatus Kaiserbacteria.

**Figure 3.**
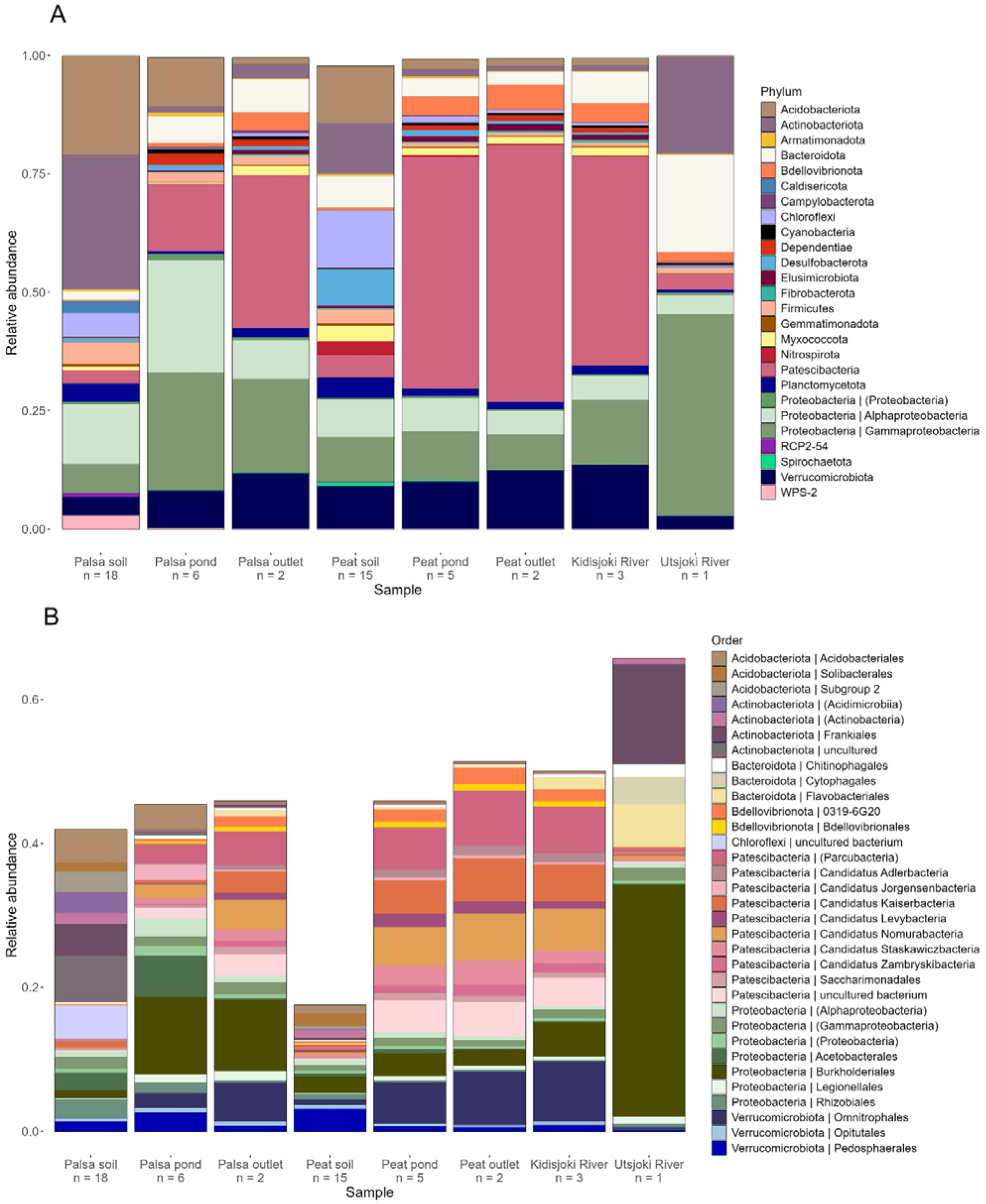
Bacterial community composition of the study site in (a) phylum level (relative abundance ≥ 0.01% shown) and (b) order level (relative abundance ≥ 0.38% shown). Number of replicates from each site is indicated below the site label in x-axis and reported relative abundances are mean values of the replicates.

#### Archaeal community composition

Archaeal communities consisted of phyla Nanoarchaeota, Micrarchaeota, Halobacterota and Crenarchaeota (Figure 4). Peat soils had highest relative abundance of archaeal 16S rRNA genes with approximately 4% of total relative abundance. The archaeal communities in palsa soils consisted of orders Group 1.1c and Methanomicrobiales, while in palsa ponds Methanobacteriales and Micrarchaeales were most abundant. Peat soil archaeal community was composed of Methanomicrobiales, Methanobacteriales and Methanosarciniales. Woesearchaeales were present in all sites, and were abundant in palsa outlet, peat soil, peat ponds, peat outlets and Kidisjoki River. Micrarchaeles were also abundant in all water samples except Utsjoki River.

**Figure 4.**
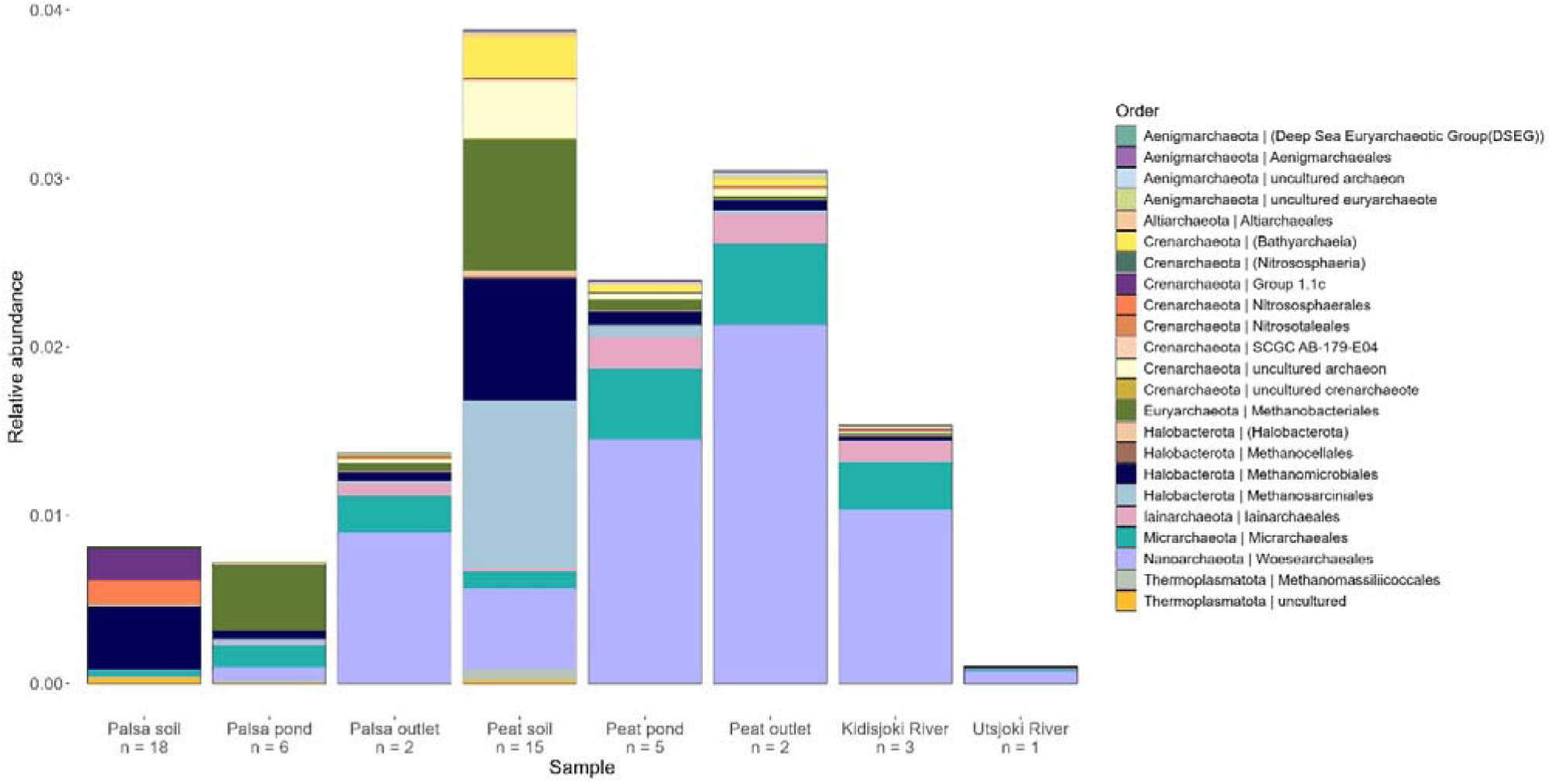
Archaeal community composition in order level of the Kidisjoki River catchment sampling sites. Number of replicate samples indicated below the site label in x-axis and relative abundances are reported as mean values of the replicates.

#### Microbial community variability

PCoA shows clear division of the microbial communities among sampled sites, with x-axis explaining 35.9 % of variation and y-axis 20.7% (Figure 5a). Palsa ponds, palsa soils and peat soils formed three distinct groups. Permafrost soil from site 26 and Utsjoki River grouped together with palsa ponds. The Kidisjoki River, outlet sites and peat ponds clustered together, forming a fourth distinct group. Palsa ponds clustered separately from other sites also in PCoA with only the water samples (Figure 5b). The Utsjoki River sample seemed to group closer to palsa ponds than to the rest of the catchment, likely due to Patescibacteria and Proteobacteria relative abundance being similar range in both sites. Peat and palsa outlets were grouped together with the rest of the water samples, unlike in the PCA of biogeochemical properties, where they formed a slightly distinct group (Figure 2).

**Figure 5.**
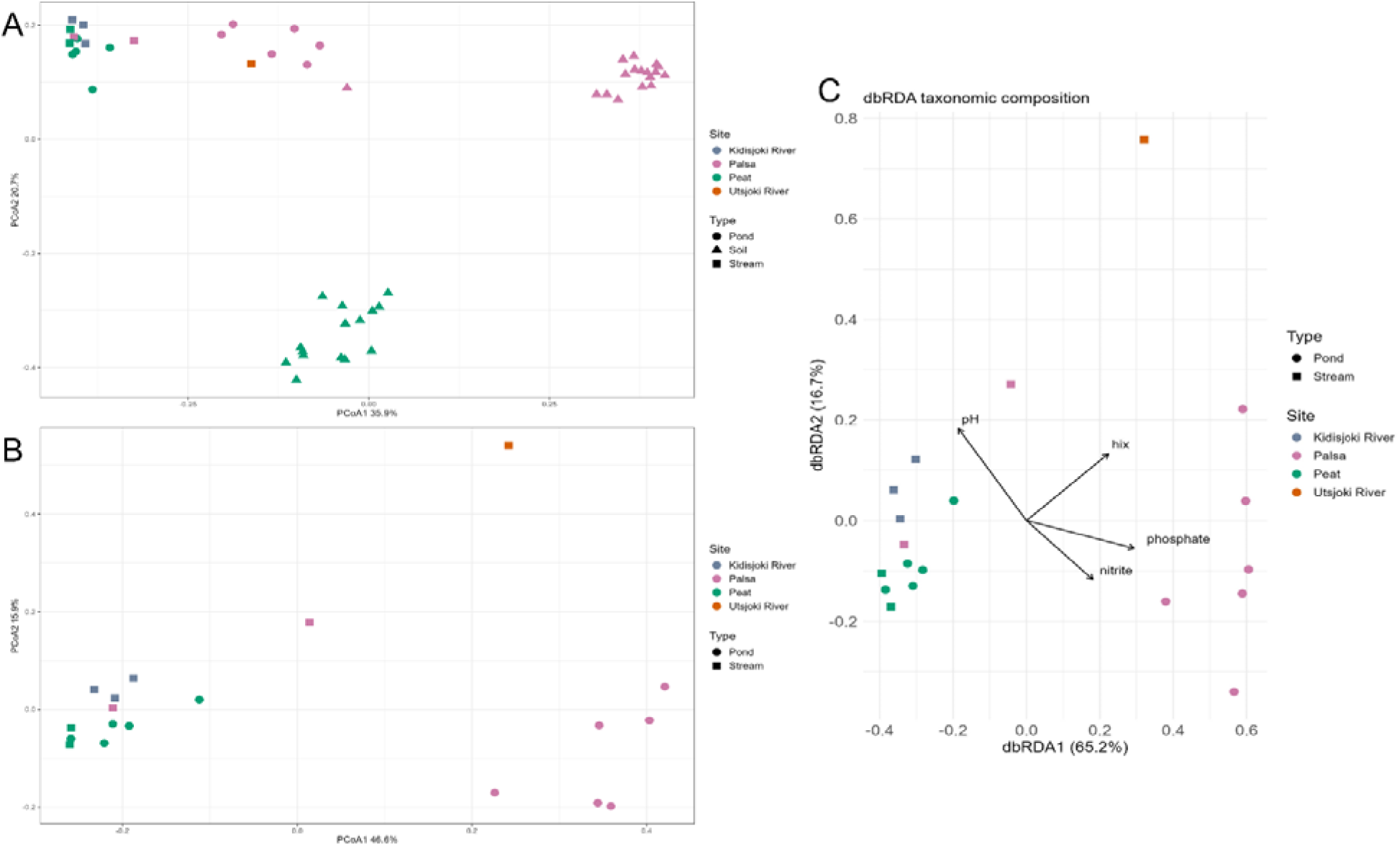
Principal Coordinate Analysis to detect variation between microbial communities of relative abundance in genus level in (a) all sampling sites and (b) sampled freshwater sites. Palsa and peat soils formed separate groups from water samples. Notably palsa ponds formed a distinct group from rest of the water sites. Sample type is indicated by shape and sample site by colour. Distance-based redundancy analysis indicated pH, hix, phosphate and nitrite to influence the microbial community composition (c).

With the dbRDA analysis, the model explained 37.2% of the total variation, with pH, nitrite, phosphate and hix as predictors (F = 2.072, p < 0.03). Permutation tests showed a significant contribution of phosphate (F = 3.936, p< 0.01), pH (F =3.120, p < 0.03) hix (F = 3.007, p < 0.03) and nitrite (F = 3.039, p < 0.03). Phosphate explained 14.5% of the variation, hix 11.1%, pH 11.5% and nitrite 11.2%.

### DOM degradation experiment

The local microbial community in the palsa pond was more active compared to any other site, as seen by higher bacterial respiration rate (Supplementary Figure 8a) and growth (Supplementary Figure 8b, Supplementary Table 2). Microbial transformation of DOM resulted in an initial increase in CDOM absorption, especially in palsa ponds (Supplementary Figure 9, Supplementary Table 2). At other sites, aCDOM_(254)_ generally decreased while absorption coefficients at higher wavelengths increased. Spectral slopes changed negligibly (Supplementary Figure 10, Supplementary Table 2). Fluorescence peaks generally increased throughout the experiment, except at palsa ponds, where they abruptly dropped between the 2^nd^ and 3^rd^ subsamplings (Supplementary Figure 11, Supplementary Table 2). At the end of the degradation experiment most CDOM and FDOM parameters had either decreased from initial values or increased very little, except for HIX which increased at each site. With a few exceptions nutrient concentrations did not change considerably during the experiment (Supplementary Figure 12, Supplementary Table 2). The exceptions are ammonium, which increased in the Utsjoki River and decreased in the palsa pond, and silicate, which increased in every site except the palsa pond.

The changes in the CDOM absorption coefficients during the degradation experiment in the palsa pond were modest compared to other sites, when accounting for the high differences in starting conditions. This suggests that the high bacterial respiration and growth could more strongly be sustained by the non-coloured DOM in the ponds.

DOC consumption in the degradation experiment was estimated using three methods for predicting DOC concentration from CDOM absorption coefficients [77–79]. Predicted DOC concentrations are highly variable, and especially the results for the palsa pond should be assessed critically, because the conversion methods were not created using such high absorption coefficient values. Yet the predicted DOC degradation of other sites was much lower than in the palsa pond, generally less than 20 µmol C L^-1^ consumed during the total duration of the experiment (Supplementary Table 2).

Non-filtered samples showed that grazers limit bacterial growth at the selected control sites (Supplementary Figure 8c). In these samples CDOM absorption coefficients decrease already during the first measurement interval (Supplementary Table 3), showing the role of bacteria in producing CDOM.

### DOC export rates

Export rates of DOC were similar in both palsa and peat outlets (Table 3). DOC export rate increased downstream reflecting DOC contribution of sub-catchments and surface flow.

**Table 3.** Concentration and export rates of dissolved organic carbon (DOC) of the catchment sampling locations estimated based on DOC and flow rate measurements in early June. Utsjoki River export rate is calculated from DOC concentration measured in this study and day specific flow rate from Finnish Environment Institute [50].

| Site name | DOC<br>g m<sup>-3</sup> | Flow rate<br>m<sup>3</sup> s<sup>-1</sup> | DOC export<br>kg C day<sup>-1</sup> |
| --- | --- | --- | --- |
| Palsa outlet 2 | 4.04 | 0.016 | 5.48 |
| Peat outlet 2 | 4.14 | 0.015 | 5.47 |
| Kidisjoki River 1 | 2.74 | 0.087 | 20.47 |
| Kidisjoki River 2 | 2.71 | 0.275 | 64.42 |
| Kidisjoki River 3 | 2.58 | 0.301 | 67.03 |
| Utsjoki River | 2.85 | 17.74 | 4 366 |

## Discussion

The main aim of the study was to identify how DOM quantity and quality was affected by peatland and degrading permafrost in a small sub-arctic catchment and how DOM properties affect the microbial community composition. We were able to capture a unique time towards the end of the spring freshet, with the catchment being hydrologically connected. During our sampling campaign palsa ponds were evaporating or infiltrating into the soils rapidly and likely had disappeared within days following our sampling.

### Dissolved organic matter from palsa mire is not visible along the catchment

Our results show high DOC concentrations and distinct DOM properties in the ponds of degrading palsa mire compared to the peatland site and the rest of the catchment. Palsa ponds were also characterized by high DOM processing and bacterial cell numbers, highlighting the role of these small water bodies in the processing of soil derived organic matter and nitrogen [14,80]. Physicochemical properties of the palsa pond were also distinct from the other sites, even the headwater stream draining directly from the palsa mire. Carbon export rate from peatland was analogous to the palsa export rate. However, the peat ponds had bacterial community reflective to the rest of the catchment, unlike the permafrost affected site.

DOC export dynamics and changes in hydrology are dependent on the development of a palsa mire to either a fen or a bog following permafrost degradation [13]. Fens remain more hydrologically connected year-round, whereas bogs are more dominated by the spring snowmelt [13]. In the Siberian Arctic, peat bog depressions of similar size as our sampled ponds had comparable DOC concentrations [19]. The study site had less dramatic decrease in DOC concentration from the depressions to small streams compared to our results. Higher DOC concentrations in thaw ponds than non-permafrost affected ponds have also been detected in Canadian Arctic [81], supporting our findings. However, our thaw ponds had overall slightly higher DOC values. This could be due to sampling time, as our sampling was done towards the end of the spring snowmelt and theirs in August during the base flow period.

Elevated concentrations of DOC and more labile characteristics downstream from the thaw slumps were detected in a Canadian peat plateau [82]. In our study site, DOC concentration downstream from both permafrost and palsa sites decreased drastically. Notably our palsa ponds were evaporating or infiltrating into the soils, likely leading reduced hydrological connectivity. These elevated DOC concentrations were not detected in the nearby outlet sites (Palsa outlet 1). In a similar palsa mire in Norway, streams draining the site were detected to be contrasting to the palsa thaw ponds, likely due to influence of groundwater flow and hydrological isolation of thaw ponds [14].

Our results are limited to a one-time sampling of a small catchment and thus might not be applicable for large-scale generalisation of DOC dynamics and processing in the changing permafrost environments of the Arctic. Yet similar trends in been detected in other studies [13,14,83].

Streams and ponds have been measured to be sources of CO_2_ and CH_4_ [14,22,83–85]. While greenhouse gas measurements were not in the scope of our study, in a similar palsa site with comparable DOC concentrations dissolved CH_4_ concentrations were 2,000 to 5,000 times of the atmospheric equilibrium in palsa ponds [14]. Whereas thaw ponds are larger sources of CH_4_ than stream sites, opposite is true for CO_2_ [14]. Similar CH_4_ range and greenhouse gas dynamics could be expected in our study site. Relative to their areal coverage, small water bodies are an unproportionally large carbon emitter and can affect carbon balance even at landscape scales [84]. The range of carbon lateral transfer and greenhouse gas emissions remains uncertain, due to limited wider sampling campaigns on small streams [84]. A further study including DOM degradation potential or greenhouse gas measurements along the hydrological continuum could shed light on the importance of microbial processing on CO_2_ and CH_4_ emissions.

### Dissolved organic matter properties are distinct between palsa ponds and rest of the catchment

The properties of DOM divided the catchment to two distinct groups: palsa ponds and the rest of the catchment. Palsa ponds CDOM was characterized as more refractory, with higher aromaticity (SUVA_254_) and larger molecular size (S_275-295_). The palsa ponds were also characterized by high total nitrogen and lower pH relative to the other sites. Lower pH, high DOC concentrations and refractory carbon quality are typical for emerging thaw ponds, observed in Canadian discontinuous permafrost zone [40,82] and palsa mires [13,14]. Different soil properties in peat and palsa sites could lead to variability in DOC properties. Palsa soils had high SOM, whilst peat soils were shallow with lower SOM content. The quality of organic matter varies, with palsa soils containing more degraded and recalcitrant organic matter in aerobic active layer and more labile organic matter in permafrost [86]. In peatland the anoxic state of the soil prevents rapid degradation [86].

Our results on CDOM properties agree with earlier work, where DOC from palsa mire was characterized as a poorer substrate for microbial consumption [13]. Higher SUVA values, higher C:N ratio and lower FI were detected in the palsa mire compared to fens in the Swedish Arctic [13]. The changes in hydrology from a palsa degrading into a fen resulted in higher groundwater connectivity, which could result in more labile DOC. Similar dynamics were also detected in our study sites. A similar trend in permafrost degradation leading to higher terrestrial and refractory carbon in freshwaters was detected also in the circumpolar permafrost zone [11].

Refractory DOC properties could be explained by the sampling time being towards the end of spring freshet and start of baseflow period. During spring freshet more organic matter is transferred, and microbial degradation is limited by shorter residence times and lower temperatures [87]. In a subarctic peatland stream, lower quantity and quality of organic matter for microbial conditions was detected during baseflow conditions [83]. During baseflow organic matter is likely degraded closer to the source soil [83].

### Aquatic microbial communities comprised mainly of ultra-small bacterial and archaeal taxa

Microbial communities were distinct between the palsa ponds and the rest of the catchment, reflecting the biogeochemistry of the sites. This supports our hypothesis that DOC quantity and quality, along with the environmental properties like pH, are affecting the microbial community composition. Microbial community assembly in palsa ponds was likely affected by the degrading palsa soils, which caused low pH and high input of DOC. In addition to the distinct community composition, we detected high relative abundance of ultra-small taxa in both bacterial and archaeal communities.

Stream and peat ponds bacterial communities were dominated by reads assigned to Patescibacteria 16S rRNA genes, consisting of over half of the relative abundance in the stream sites and peat ponds. While this bacterial phylum is relatively unknown, it has been detected in high abundances in e.g. groundwater [88,89] wastewaters [90] and thermokarst ponds [35,40]. Of Patescibacteria, the orders Candidatus Kaiserbacteria and Candidatus Nomurabacteria were the most abundant in the stream sites, which have been detected in Arctic thermokarst lakes [35] and groundwater [91]. Patescibacteria relative abundance was lower in the Utsjoki River than in the smaller streams and ponds. While our data had only one sampling point from the Utsjoki River, similar trend of decrease in Patescibacteria with increasing river size has been detected in a large-scale riverine study [33].

Patescibacteria lifestyle is considered host dependent, either symbiotic or parasitic, due to the reduced genome size and lack of metabolic pathways [89,92]. Patescibacteria have been detected to ferment carbon compounds to acetate, and benefitting e.g. methanogens, methanotrophs, Actinobacteria, Betaproteobacteria, Planctomycetes, and Chloroflexi [35]. Patescibacteria are facultative anaerobes but possess a genetic ability to resist oxygen [35].

Ultra-small archaea Nanoarchaeota and Micrarchaeota were also detected along the catchment. They have similar lifestyle as ultra-small bacteria, with a host attached lifestyle and an ability to degrade complex organic matter by fermentation [38,39]. Archaeal communities were distinct between the palsa ponds and the rest of the catchment, with Nanoarchaeota and order Woesearchaeales dominating in the peat ponds and the stream sites, and Methanobacteriales in the palsa ponds.

Besides Patescibacteria, the palsa pond bacterial community consisted of Acidobacteriota, Proteobacteria, Bacteroidota and Verrucomicrobiota. Similar bacterial communities have been detected in thaw ponds of the Canadian Arctic [27,40]. Palsa soil was dominated by Acidobacteriota, Actinobacteriota and Proteobacteria, similarly as earlier findings in the Stordalen palsa mire in Sweden [93]. The Stordalen fen soils were comprised of a higher number of bacterial phyla, and higher archaeal relative abundance, than the palsa soils [93], supporting our findings.

Soils are the main source of bacteria into the freshwaters, where the bacterial communities are shaped by stochastic and deterministic processes [28,29,32]. Our sampled freshwaters were highly connected to soils, yet the community composition between the soil and the adjacent pond was distinct. Since our sampling was conducted towards the end of spring freshet, microbial communities in the ponds had likely been developed over few weeks, creating distinct community structure from the soils. Stochastic processes might dominate community assembly in the stream sites and ponds immediately after spring freshet [29], whereas towards the end of freshet, selection might dominate microbial community assembly, especially in the palsa ponds, where DOC and dissolved nutrient concentrations were high.

It is likely that the peatland and Kidisjoki River were also dominated by bacterial taxa derived from snowmelt or groundwater, rather than soil input. During high flow events, selection hotspots are focused on areas with longer residence times [29]. Our sampling was towards the end of high flow snowmelt, and apart from the less connected palsa ponds, the residence time could have been too short for selection to occur, and rather the mass effect of microbial input from soils and groundwater dominated the assembly processes. While this effect might not be detected here, assembly-based methods might give more insight into the formation and occurrence of similar taxa between soil and water samples. Our study is limited to the free-living and particle associated microbes. Benthic microbial communities might display the community assembly processes along the stream network [94].

## Conclusions

The impact of degrading palsas and peatland on the DOM quality and quantity and microbial community composition was studied on a sub-arctic river catchment. As expected, terrestrial DOM loading was high from both peat and palsa soils to ponds. In the palsa ponds DOM quantity and quality, physicochemical properties and carbon processing differed from the peat ponds and the downstream catchment. Similar range of DOC export was measured from both the permafrost and the peatland sites. Microbial community composition reflected the DOM quality and quantity, alongside with other environmental variables, with the taxonomic composition in the palsa ponds being distinct from the rest of the sites. The relative abundance of ultra-small Patescibacteria was high along the whole catchment. Although limited to one sampling period our results highlight that degrading palsas may not be a continuous seasonal source of microbes and organic carbon to a wider fluvial system as is frequently postulated.

## Data availability

DNA sequences have been deposited in ENA under project number PRJEB85624. Biogeochemical data is available in Pangaea (https://doi.pangaea.de/10.1594/PANGAEA.981156).

## CRediT authorship contribution statement

NT: Conceptualization; funding acquisition; field work; data curation; formal analysis; investigation; visualization; writing—original draft; writing—review and editing. SE: Conceptualization; field work; data curation; formal analysis; investigation; visualization; writing—review and editing. HK: Conceptualization; supervision; investigation; resources; writing – review & editing. JH: Conceptualization; supervision; investigation; resources; writing – review & editing. DNT: Conceptualization; project administration; field work; investigation; resources; supervision; writing – review & editing.

## Acknowledgements

We thank the funders of this study, Kone Foundation (grant number: 202204330) and Emil Aaltonen Foundation (grant number: 220226). JH acknowledges the Research Council of Finland (354462). We are grateful to the help of the staff at the Kevo Subarctic Research Institute of the University of Turku.

Metsähallitus (Parks & Wildlife, Finland) is acknowledged for sampling permit. We thank Janina Kariluoto for her assistance during the field campaign and with the laboratory work. We acknowledge DNA Sequencing and Genomics Laboratory (supported by HiLIFE and Biocenter Finland funding), Institute of Biotechnology, University of Helsinki for sequencing. We acknowledge CSC – IT Center for Science, Finland for the provision of computational resources. The study utilized marine research infrastructure at the Finnish Environment Institute as a part of the national FINMARI RI consortium. The flow cytometry analysis was performed at the HiLife Flow Cytometry Unit, University of Helsinki.

