## Supplementary for "Carbon and microbes in a degrading palsa mire are distinct from peatland and a wider connected sub-Arctic fluvial system"

Tuomela, Nea^1*^; Elovaara, Samu^2^; Hultman, Jenni^3^; Kaartokallio, Hermanni^2^; Thomas, David N.^1^

^1^ Faculty of Biological and Environmental Sciences, University of Helsinki, Helsinki, Finland

^2^ Finnish Environment Institute (SYKE), Helsinki, Finland

^3^ Natural Resources Institute Finland (LUKE), Helsinki, Finland

The following document contains supplementary material for “Carbon and microbes in a degrading palsa mire are distinct from peatland and a wider connected sub-Arctic fluvial system”.

14 Pages, 12 Figures, 1 Table

### List of Figures

**Supplementary Figure 1**. Examples of the sampled soil and water sites in palsa mire (left), and peatland site (right). In palsa site permafrost degradation was visible with collapse features.

**Supplementary Figure 2**. Bacterial community composition in phylum level based 16S rRNA gene annotations from metagenomes divided by samples site (a) palsa, (b) peat and (c) streams sites.

**Supplementary Figure 3.** Bacterial community composition in order level based 16S rRNA gene annotations from metagenomes divided by samples site (a) palsa, (b) peat and (c) streams sites.

**Supplementary Figure 4.** Archaeal community composition in order level based 16S rRNA gene annotations from metagenomes divided by samples site (a) palsa, (b) peat and (c) streams sites.

**Supplementary Figure 5.** Gating examples used in to derive data for bacterial cell number data for (a) high nucleic acid and (b) low nucleic acid cells.

**Supplementary Figure 6.** Shannon diversity index of the sampled soil and water sites.

**Supplementary Figure 7.** Screeplot used to justify the selection of principal components with the biogeochemical data.

**Supplementary Figure 8.** Microbial respiration rate (a) and high nucleic acid bacterial cell numbers (b,c) during degradation experiment

**Supplementary Figure 9.** CDOM absorption coefficients at selected wavelengths in the peat, Kidisjoki River and Utsjoki River sites (a) and palsa sites (b) during the degradation experiment.

**Supplementary Figure 10.** CDOM spectral slopes at selected wavelength ranges and the S_ratio_ in the peat, Kidisjoki River and Utsjoki River sites (a) and palsa sites (b) during the degradation experiment.

**Supplementary Figure 11.** Coble’s peaks A, C, M and T of FDOM and the biological index, the humification index and the fluorescence index in the peat, Kidisjoki River and Utsjoki River sites (a) and palsa sites (b) during the degradation experiment.

**Supplementary Figure 12.** Nutrient concentrations in the peat, Kidisjoki River and Utsjoki River sites (a) and palsa sites (b) during the degradation experiment.

### Supplementary Methods

#### Supplementary Methods 1: Cell Abundance

Bacterial cell number was analyzed using flow cytometry with LSRFortessa (BD Biosciences) at University of Helsinki Flow Cytometry Unit. Samples were fixed with paraformaldehyde (concentration 0.9%), incubated for 15 minutes in the dark and stored in -80 °C until analysis. Prior to analysis samples were thawed, filtered with strain cap tubes (pore size 35 μm, Fisher Scientific), stained with Sybr Green I nucleic acid stain (final concentration 1:10 000 Vol.) and incubated for 10 minutes in dark and room temperature. Each sample was measured in triplicate and controls with ultrapure water were run after every 10 measurements. Palsa pond samples were diluted in 1:10 ratio with ultrapure water (MQ). CountBright™ Absolute Counting Beads (Thermo Fisher Scientific) were used to calculate the concentration of cells per millilitre. Bacteria were detected and counted by plotting green fluorescence against side scatter (Supplementary Figure 5) with FlowJo software v.10.10.0 (BD Biosciences).

### Supplementary Figures

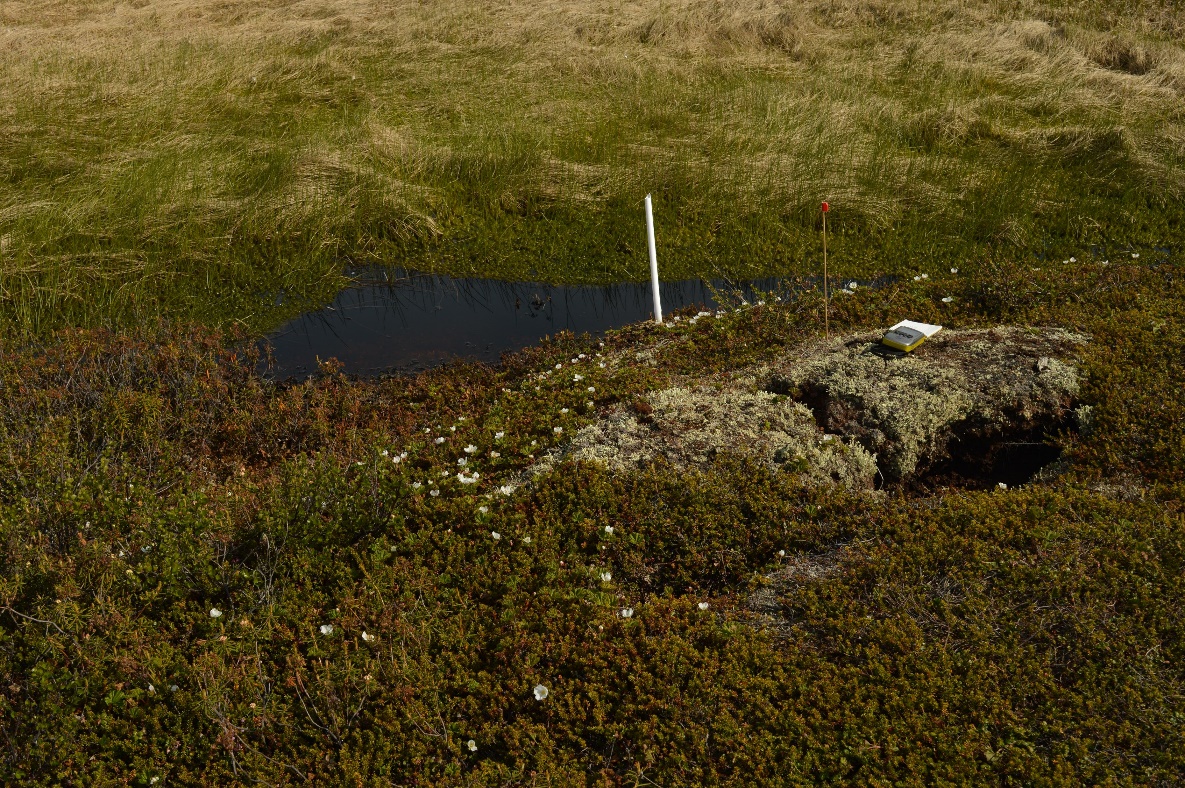

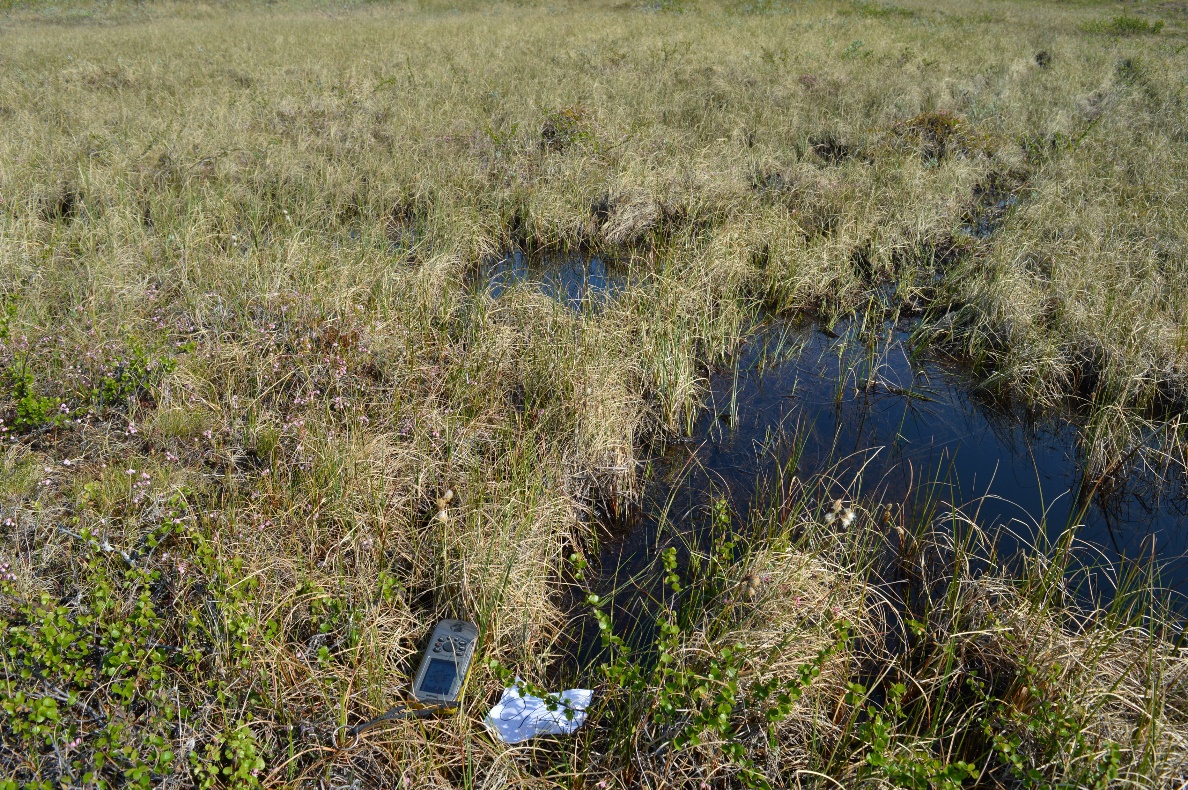

Supplementary Figure 1. Examples of the sampled soil and water sites in palsa mire (left), and peatland site (right). In palsa site permafrost degradation was visible with collapse features.

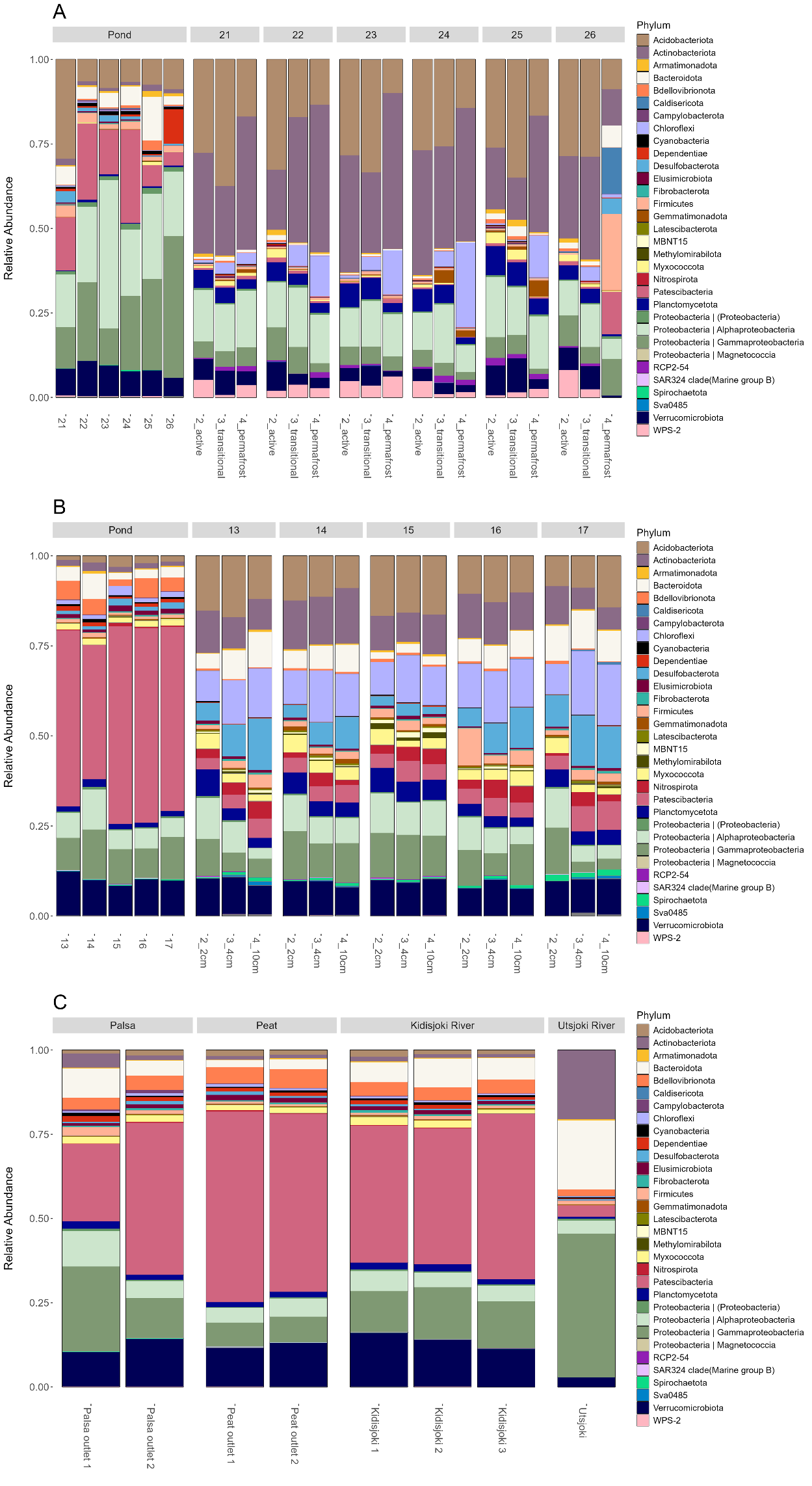

Supplementary Figure 2. Bacterial community composition in phylum level for each individual sample in (a) palsa site, (b) peat site and (c) stream sites. For palsa soils, active layer refers to the dry peat layer (20-25cm below ground), transitional to the depth of freezing in the soil (23-33cm) and permafrost the frozen soil (49-59cm). In stream sites outlet 1 is left bar and outlet 2 is the right one. Relative abundance lower than 0.01% are filtered out.

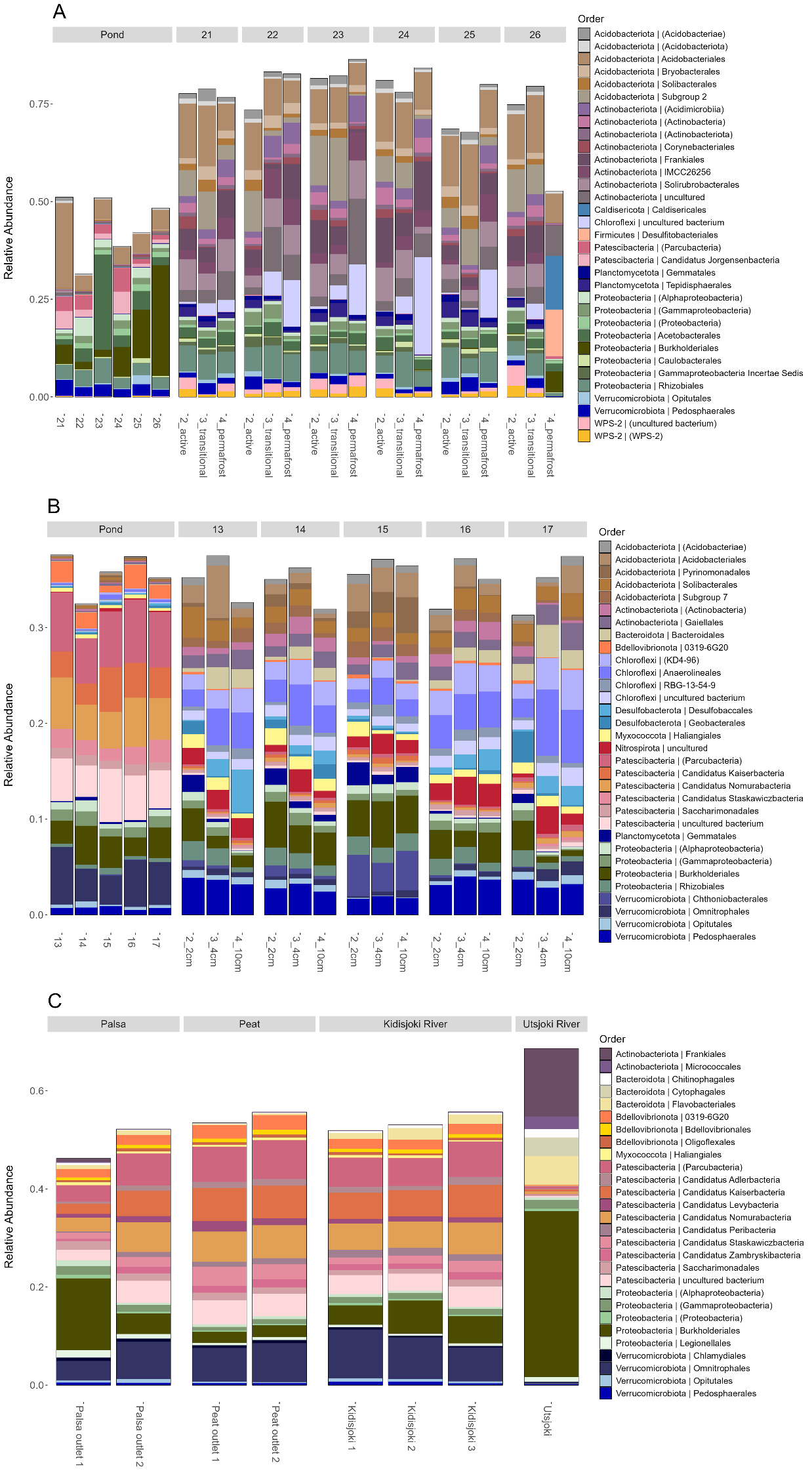

Supplementary Figure 3. Bacterial community composition in order level for each individual samples (a) palsa site, (b) peat site and (c) stream sites. In stream sites outlet 1 is left bar and outlet 2 is the right one. Relative abundance lower than 0.38% are filtered out.

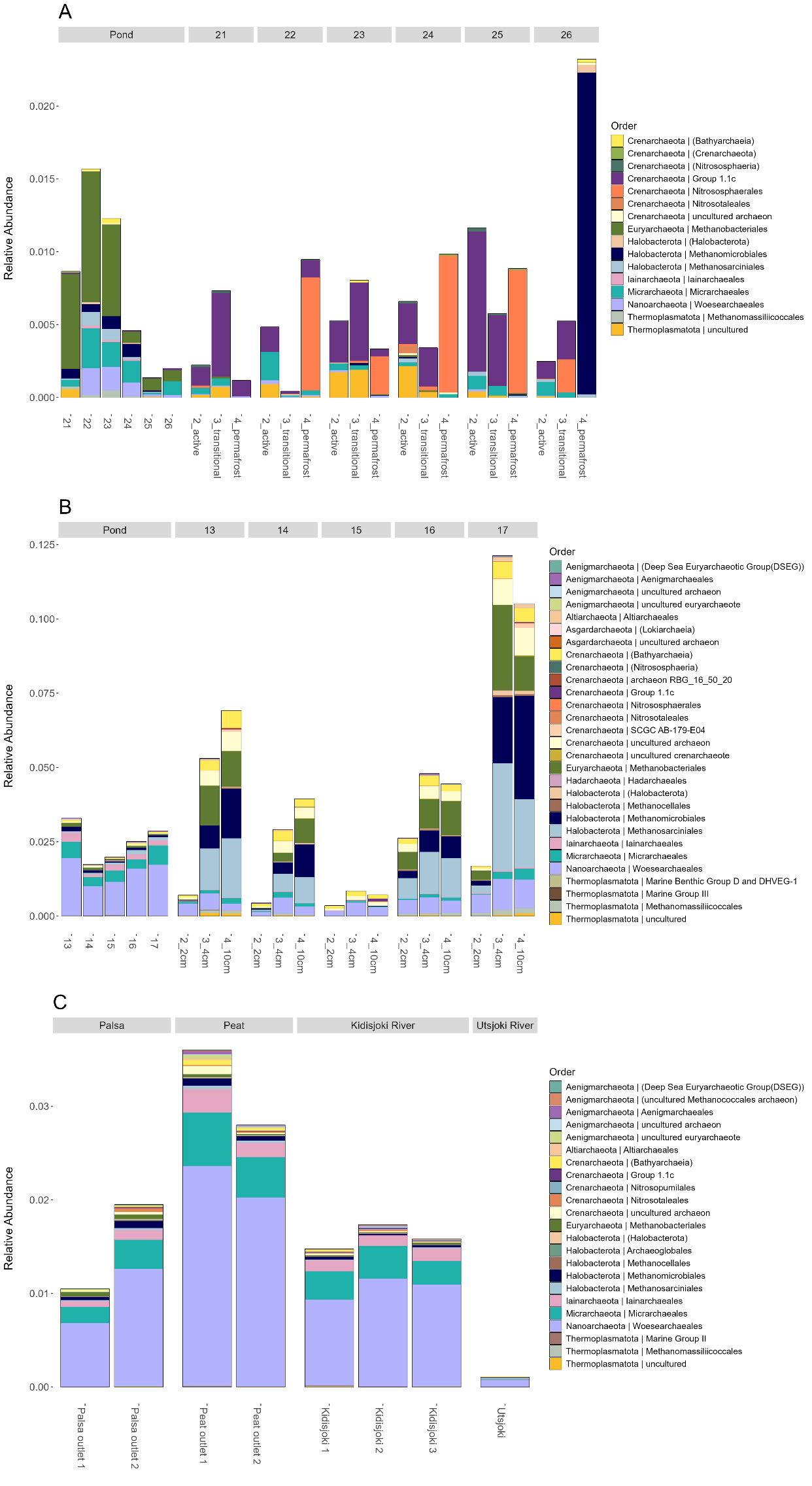

Supplementary Figure 4. Archaeal communities in order level for each sample in (a) palsa site, (b) peat site and (c) stream sites. In stream sites outlet 1 is left bar and outlet 2 is the right one.

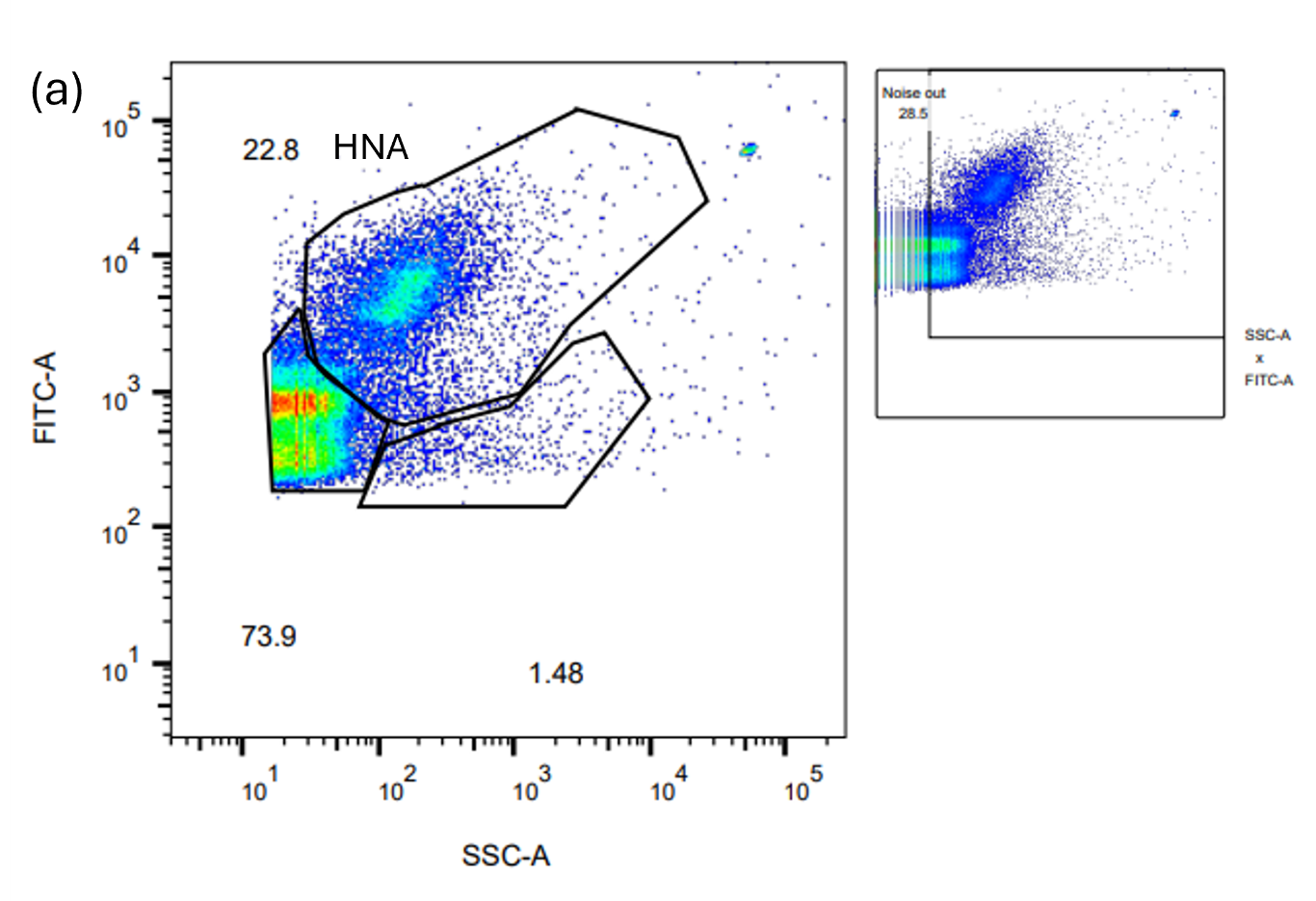

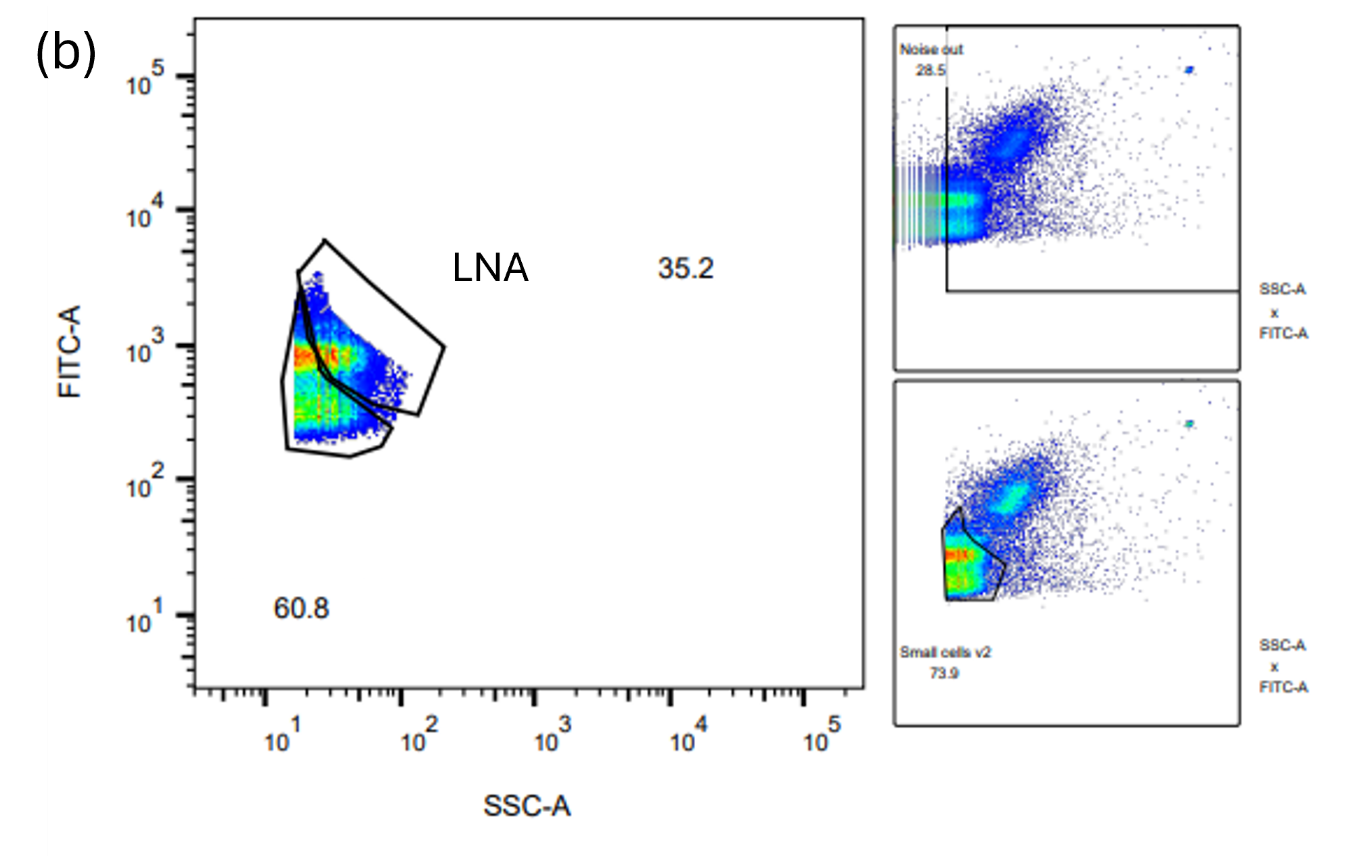

Supplementary Figure 5. Examples of the gating approach used for cell enumeration of (a) HNA cells and (b) LNA in diluted palsa pond samples

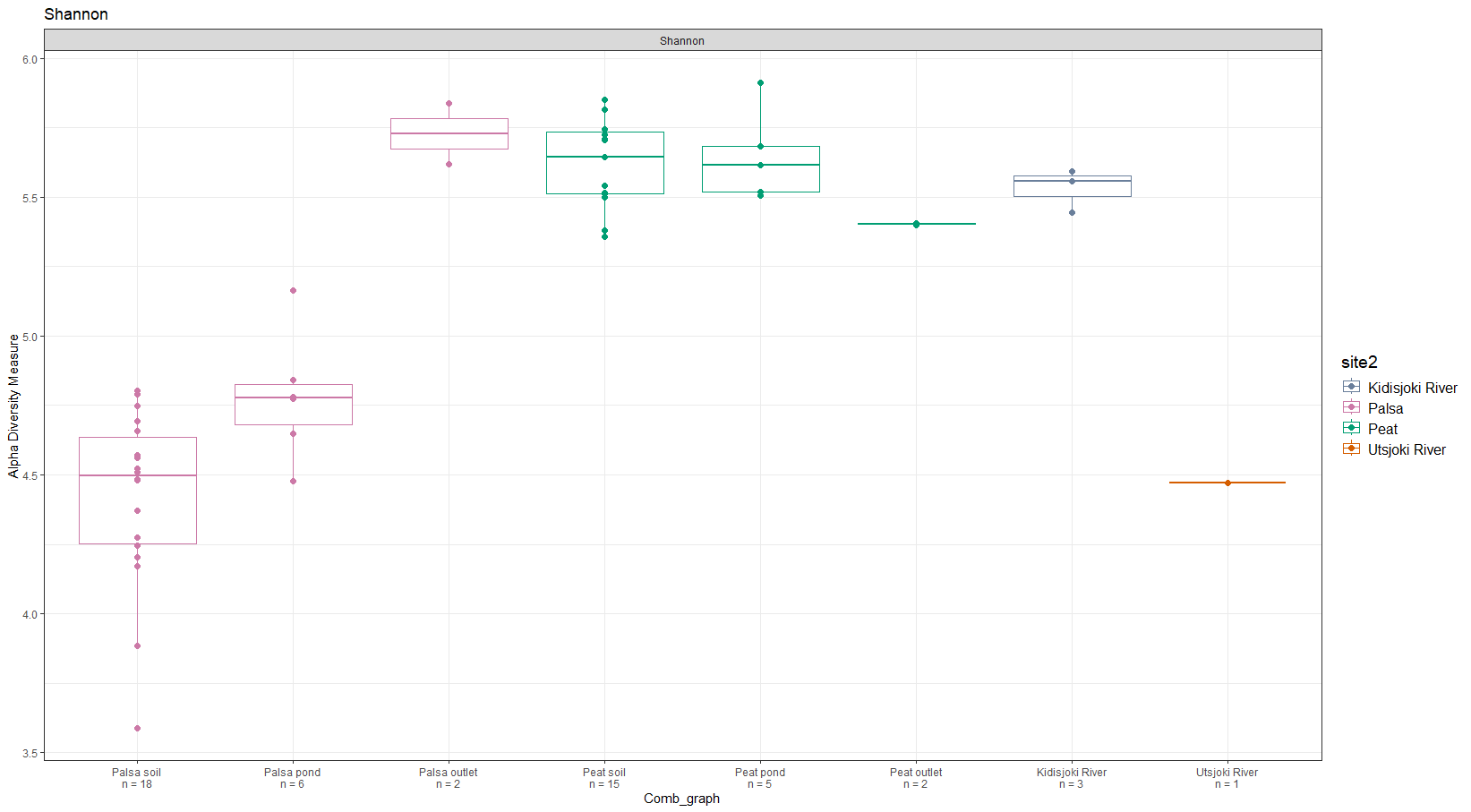

Supplementary Figure 6. Shannon diversity index of the 16S rRNA gene annotations from metagenomes. Total number of samples indicated in x-axis below the sample name.

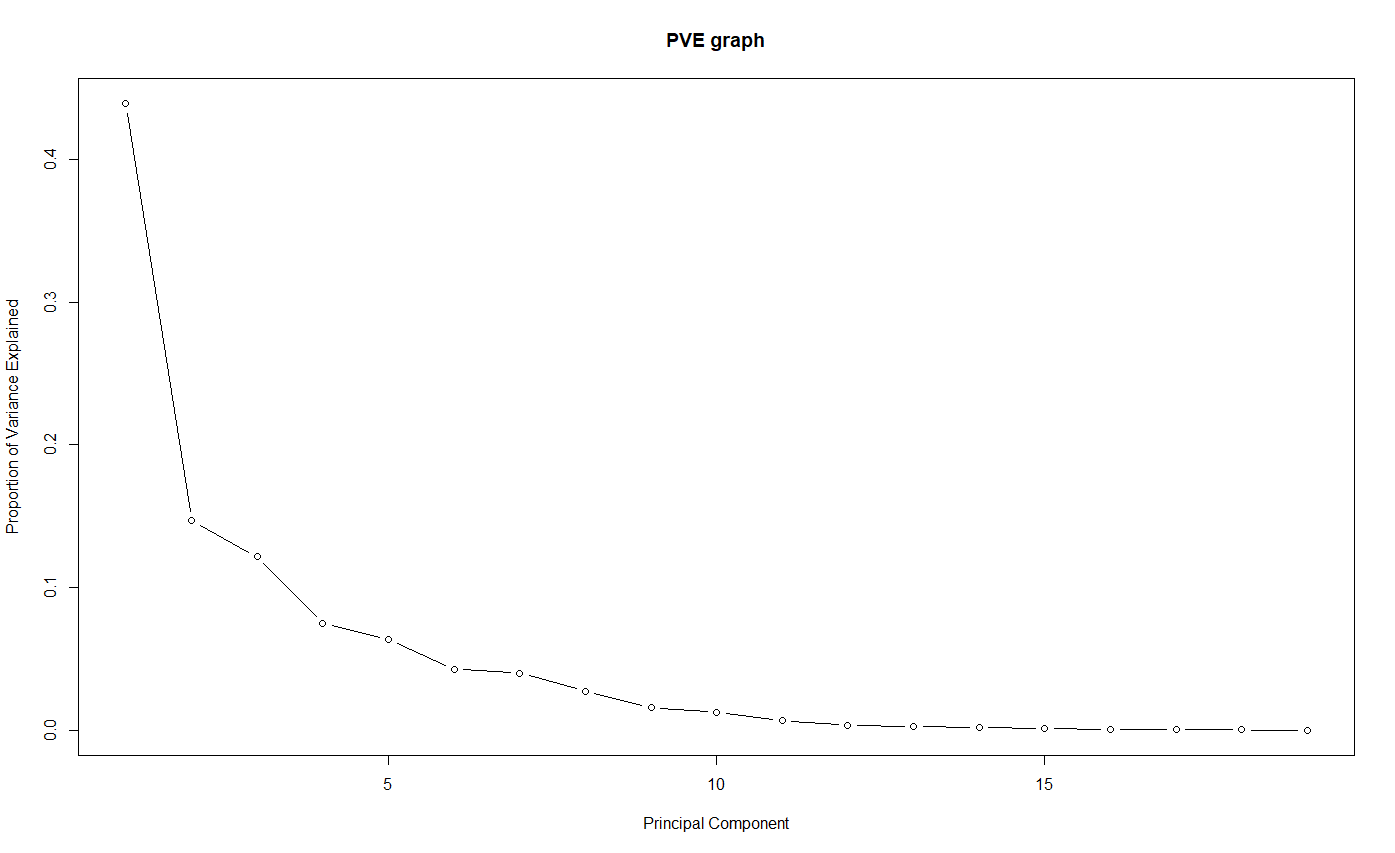

Supplementary Figure 7. Screeplot was used to justify the selection of principal components with the biogeochemical data.

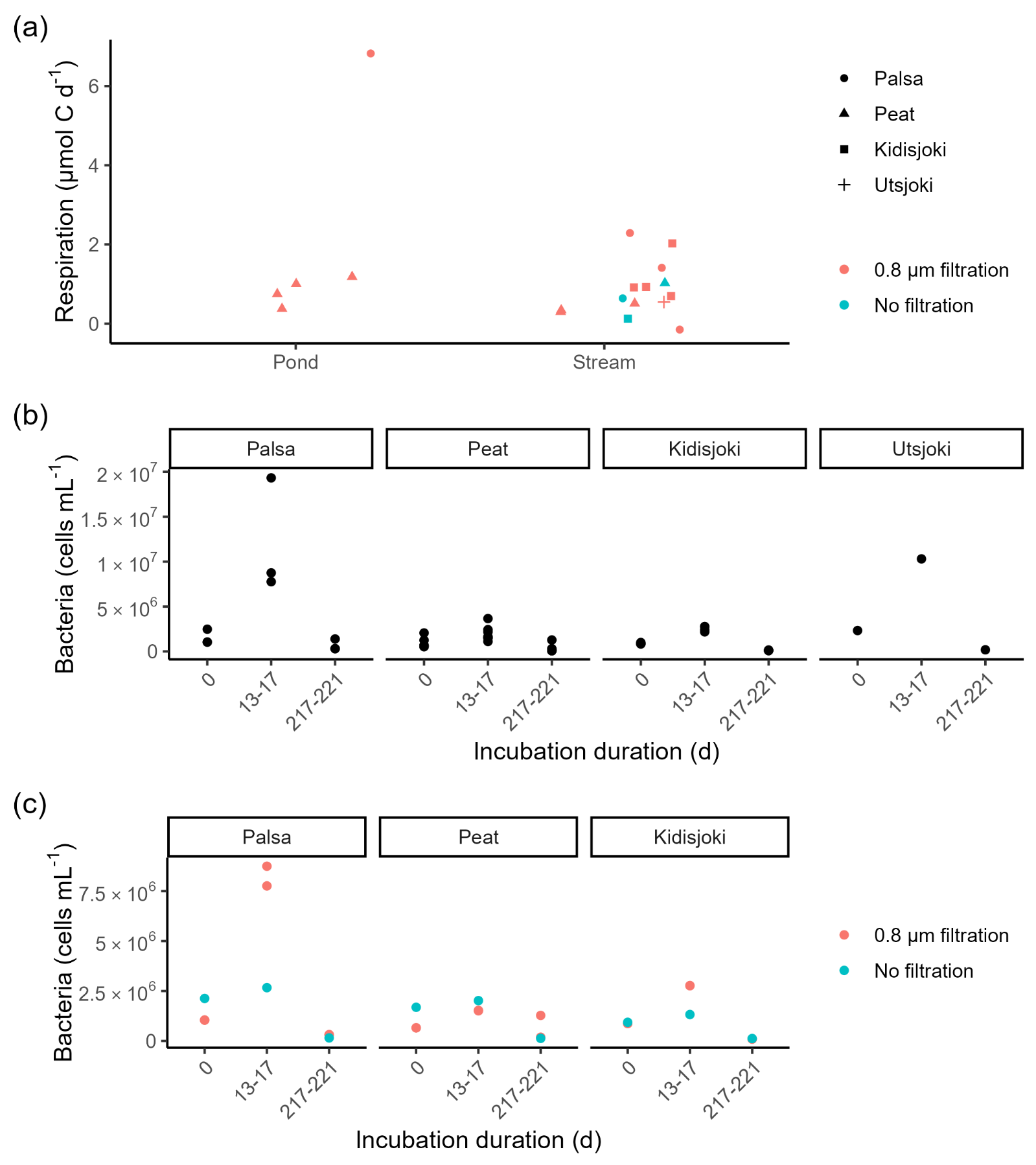

Supplementary Figure 8: Microbial respiration rate (µmol C L^1-^ d^-1^) during the first 13-17 days of the degradation experiment (a). Abundance of HNA bacteria in all filtered degradation experiment samples except the palsa pond, which was removed because of much higher HNA abundance (b). Abundance of HNA bacteria in the stream sites where the non-filtered full community controls where collected (c).

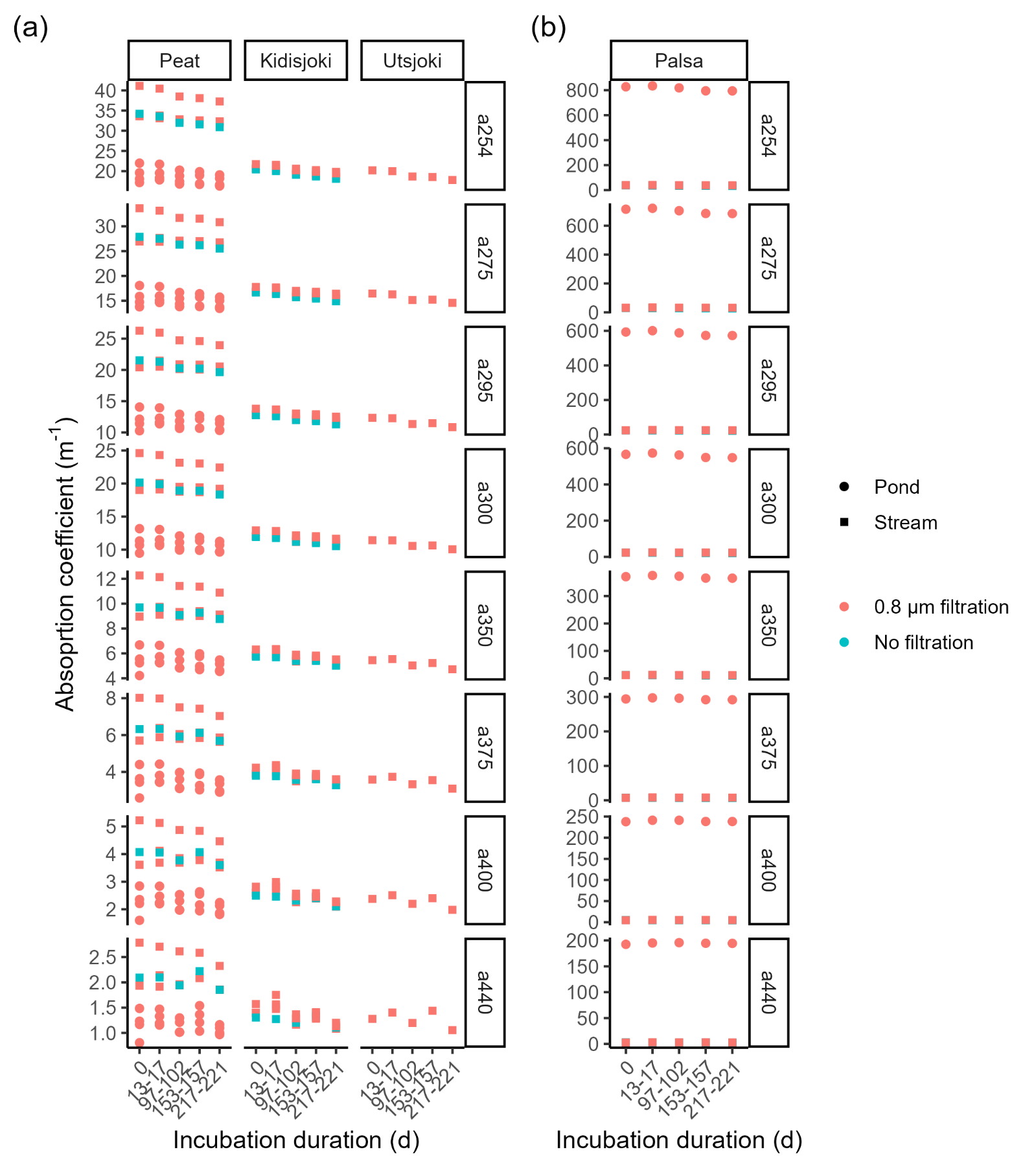

Supplementary Figure 9: CDOM absorption coefficients at selected wavelengths in the peat, Kidisjoki River and Utsjoki River sites (a) and palsa sites (b) during the degradation experiment.

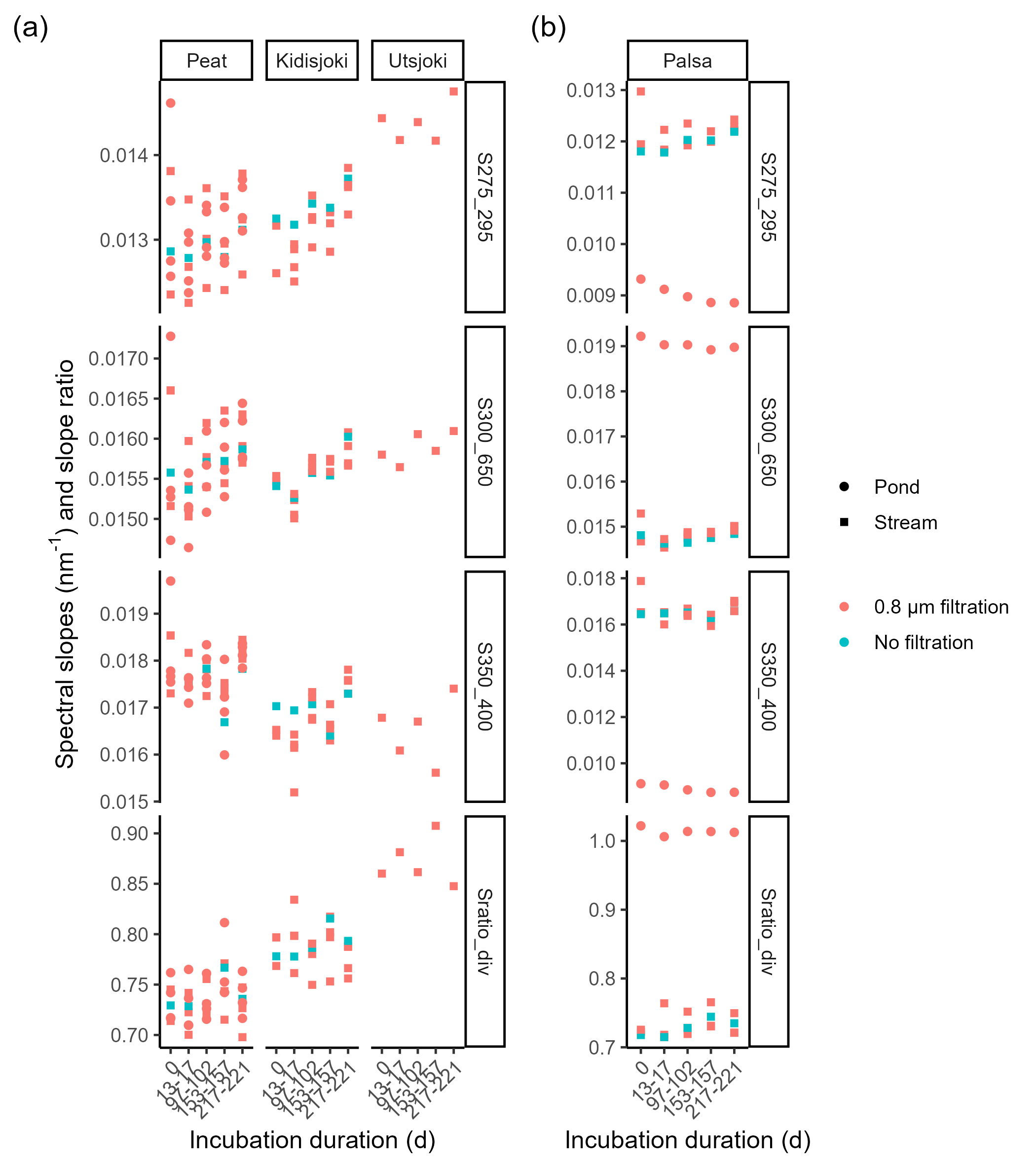

Supplementary Figure 10: CDOM spectral slopes at selected wavelength ranges and the S_ratio_ in the peat, Kidisjoki River and Utsjoki River sites (a) and palsa sites (b) during the degradation experiment.

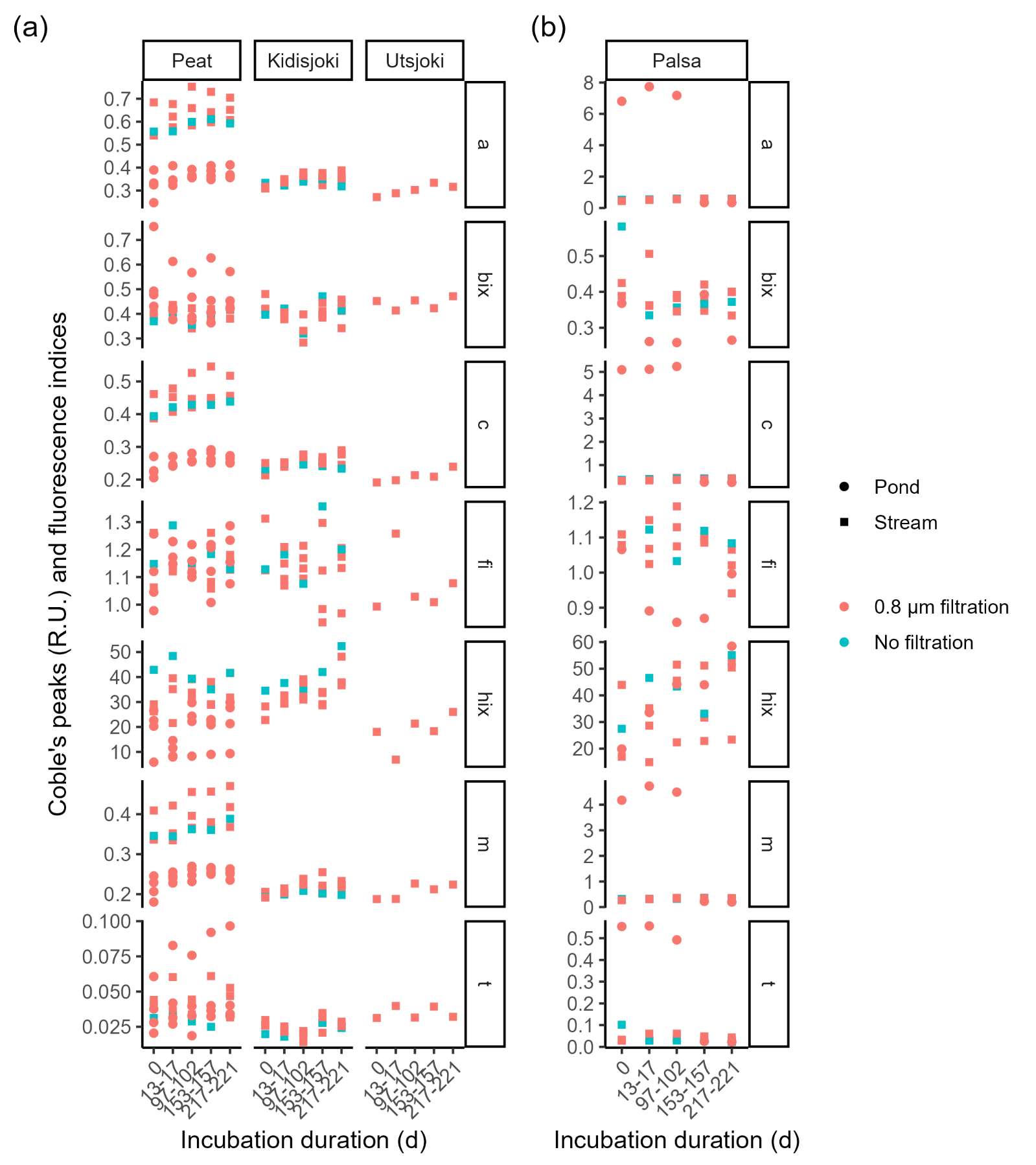

Supplementary Figure 11: Coble’s peaks A, C, M and T of FDOM and the biological index (bix), the humification index (hix) and the fluorescence index (fi) in the peat, Kidisjoki River and Utsjoki River sites (a) and palsa sites (b) during the degradation experiment.

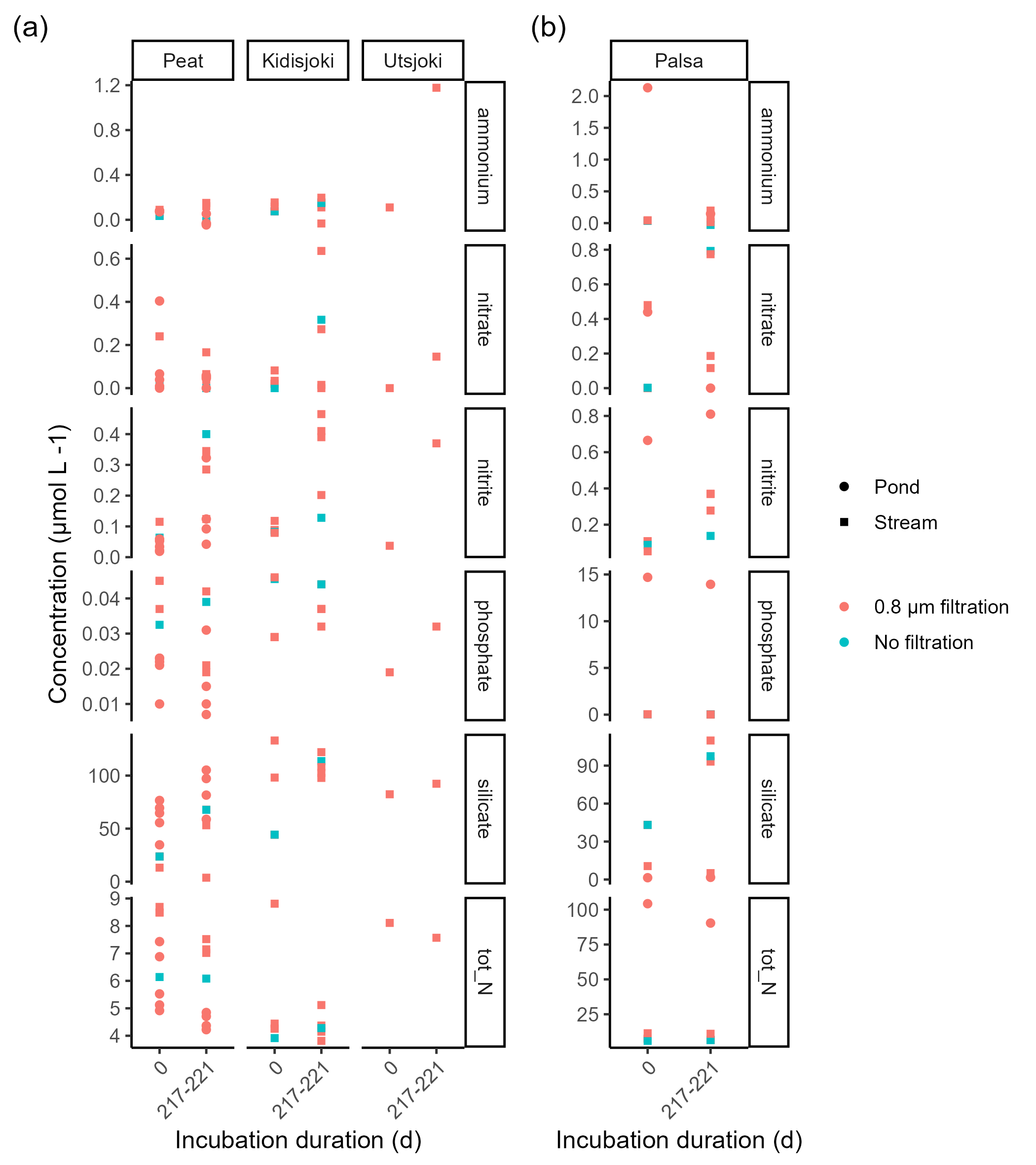

Supplementary Figure 12: Nutrient concentrations in the peat, Kidisjoki River and Utsjoki River sites (a) and palsa sites (b) during the degradation experiment.

### Supplementary Tables

Supplementary Table 1. Filtered water volumes for the microbial community composition. Each sample was filtered sequentially with 0.45µm and 0.1µm pore sized filters. DNA extraction was performed individually and extracted DNA was combined with same ratio as the filtered volumes prior to sequencing.

| Sample | 0.45 µm | 0.1 µm |
| --- | --- | --- |
|  | (ml) | (ml) |
| 1 | 1000 | 500 |
| 4 | 1060 | 500 |
| 5 | 1400 | 616 |
| 10 | 560 | 500 |
| 13 | 630 | 500 |
| 14 | 387 | 387 |
| 15 | 470 | 282 |
| 16 | 366 | 366 |
| 17 | 262 | 262 |
| 18 | 500 | 498 |
| 21 | 66 | 66 |
| 22 | 64 | 64 |
| 23 | 40 | 40 |
| 24 | 36 | 36 |
| 25 | 28 | 26 |
| 26 | 130 | 84 |
| 27 | 398 | 402 |
| 28 | 340 | 398 |
| 30 | 500 | 330 |
